# Towards large-scale museomics projects: a cost-effective and high-throughput extraction method for obtaining historical DNA from museum insect specimens

**DOI:** 10.1101/2024.09.11.612573

**Authors:** Anna J. Holmquist, Holly Tavris, Grace Kim, Lauren A. Esposito, Brian L. Fisher, Athena Lam

## Abstract

Natural history collections serve as invaluable repositories of biodiversity data. Large-scale genomic analysis would greatly expand the utility and accessibility of museum collections but the high cost and time-intensive nature of genomic methods limit such projects, particularly for invertebrate specimens. This paper presents an innovative, cost-effective and high-throughput approach to extracting genomic DNA from diverse insect specimens using single-phase reverse immobilization (SPRI) beads. We optimized PEG-8000 and NaCl concentrations to balance DNA yield and purity, reducing reagent cost to 6-11 cents per sample. Our method was validated against three widely used extraction protocols, and showed comparable DNA yield and amplification success to the widely used Qiagen DNeasy kit. We successfully applied the protocol in a high-throughput manner, extracting DNA from 3,786 insect specimens across a broad range of ages, taxonomies, and tissue types. A detailed protocol is provided to facilitate the adoption of the method by other researchers. By improving one of the most crucial steps in any molecular project, this SPRI bead-based DNA extraction approach has significant potential for enabling large-scale museomics projects, thereby increasing the utility of historical collections for biodiversity research and conservation efforts.

## Introduction

Museum collections are underutilized resources in the biodiversity sciences, containing hundreds of thousands of specimens with reliable taxonomic identifications and other associated ecological and spatiotemporal data. Recent technological advancements have enabled large-scale digitization efforts as well as the generation of genomic data, greatly increasing the value and accessibility of museum collections. Molecular methods including restriction digest, whole genome sequencing, UCE capture, and amplicon sequencing have become common approaches for obtaining information from historical DNA. However, these approaches are cost- and time-prohibitive, particularly for invertebrate collections which are typically the largest and most diverse component of a museum’s collection. To address these challenges, there is a need for high-throughput, low-cost protocols that produce archival-quality DNA, allowing researchers to unlock more of the valuable information housed in these collections.

DNA barcoding methods have become the most common approach to obtaining genetic information from thousands of specimens in a high-throughput and cost-effective manner (Hebert, Ratnasingham, and De Waard 2003; Fišer Pečnikar and Buzan 2014). Barcoding becomes especially powerful when used for metabarcoding, in which samples such as bulk biotic community collections, gut contents, or environmental DNA are sequenced. Such approaches have been used to quantify the extent of biodiversity in many understudied groups, including benthic macroinvertebrates (Hajibabaei et al. 2012) and zooplankton (Zhao et al. 2021), as well as to quantify the effects of disturbance on biotic communities (Holmquist et al. 2024). However, incomplete DNA reference databases limit the power of metabarcoding studies by preventing the assignment of taxonomic identities, requiring molecular operational taxonomic units (mOTUs) to replace taxon identity in community analyses. To gain essential information on communities through metabarcoding, robust DNA reference libraries are needed that link a barcode sequence to an expertly verified taxonomic identity.

Museum collections are powerful resources for constructing reference libraries, housing the holotype and paratype specimens used in taxonomic descriptions as well as thousands of other identified specimens. Previous studies have attempted to generate large numbers of DNA barcodes from museum collections. One of the largest projects sequenced over 41,000 Lepidoptera specimens from the Australian National Insect Collection (Hebert et al. 2013). A more recent study using the Diptera collection at the Smithsonian Institution generated 867 sequences from 941 samples using next-generation sequencing techniques (Levesque-Beaudin et al. 2022). While these and other studies have made immense strides in the processing of a large number of museum specimens, there are still barriers that need addressing, particularly reducing costs, increasing taxonomic breadth, and improving protocol accessibility to enable use by the broader scientific community.

The first step in any museomics project is the isolation of the highest quality DNA possible. Historical DNA is expected to be highly degraded with extensive sheering, depurination, and deamination (Raxworthy and Smith 2021). Extract yields are additionally expected to be low, making the extraction step incredibly important for downstream applications. While not targeted for historical DNA, Qiagen DNeasy kits (Qiagen, Hilden, Germany) are commonly used for extracting DNA from specimens in museum collections. A 4 x 96 DNeasy Blood & Tissue kit costs roughly $1,800, corresponding to about $4 per sample for extractions. Another popular and more cost-effective approach for museum specimens is the use of silica membrane plates for DNA purification following lysis (Hebert et al. 2013; Ivanova, Dewaard, and Hebert 2006). The PALL-brand filtration plates (Danaher Corporation, Washington, D.C., USA) commonly used for this method have a per-sample cost that can be as low as 54 cents per sample. However, this price covers only the filtration plates; while this method remains much cheaper than DNeasy kits, it is still prohibitively expensive for large-scale projects when the costs of necessary reagents and consumables are included.

Single-phase reverse immobilization (SPRI) bead-based approaches are one potential method to generate high-quality DNA extracts from museum samples at a low price point. While most commonly used for size selection and purification of sequencing libraries, bead-based methods can be highly effective in DNA isolation (Liu et al. 2023). SPRI methods use paramagnetic beads coated with carboxyl groups that bind nucleic acids when in a solution of salt and polyethylene glycol (PEG). The presence of salt controls the ionic strength of the solution, facilitating the binding of nucleic acids to the surface of the beads. PEG acts as a molecular crowding agent and reduces the solubility of nucleic acids, effectively precipitating them onto the beads by increasing the likelihood of interaction between the carboxyl groups and the nucleic acids’ phosphate backbones. The beads are separated from the solution with a magnetic field, immobilizing the nucleic acids, which then allows the removal of the remaining supernatant containing unwanted proteins, salts, and other contaminants. The nucleic acids are then unbound from the surface of the beads by resuspension and the resulting elute contains high-quality, archivable DNA. Bead cleaning procedures require fewer steps than other protocols, reducing the risk of additional DNA shearing that can occur through multiple rounds of centrifugation and pipetting - an important consideration when working with historical DNA. It is additionally fast and cost-effective when performed using buffers prepared in-house, making it an attractive option for large-scale studies.

The primary question in our study was: can a SPRI bead-based extraction protocol work as an effective high-throughput and low-cost method for obtaining DNA from different kinds of insect specimens from museum collections? To answer this question, we optimized bead solutions and protocol components and then compared the quality of DNA extracts from our approach against other extraction protocols. We additionally tested the high-throughput potential of the protocol on thousands of insect specimens from different orders, preserved using different manners, and collected over different time intervals. We found our extraction method to be effective in generating high-quality and amplifiable DNA from all manners of samples, with only one hour of direct labor following cell lysis and at a reagent cost of 6–11 cents per sample depending on sample type.

## Materials and Methods

### Experiment 1

Our first goal was to optimize a solution that minimized the ratio of beads necessary while retaining the ability to bind small DNA fragments. Our starting solution was based on the bead recipe published in Rohland and Reich 2012, consisting of 0.1% Sera-Mag Magnetic Speed-beads (Danaher Corporation), 18% PEG-8000, 1 M NaCl, 1 mM EDTA, 10 mM Tris-HCl, and 0.05% Tween. We began optimization by altering the concentration of PEG-8000. We hypothesized that we would be able to reduce the volume of beads needed while retaining small DNA fragments by increasing the PEG concentration. We tested PEG concentrations from 18–22% in 1% increments. We also tested a gradient of bead ratios (0.6x, 0.8x, 1x, 1.2x, 1.4x, 1.8x, 2x) paired with each PEG concentration.

Each bead solution was used to perform size selection on a diluted GeneRuler 100bp DNA ladder (Thermo Fisher Scientific, Waltham, MA, USA). 2 μL of GeneRuler was added to 18 μL of double-distilled H2O (ddH2O) to create test samples. Bead solutions were added to each sample with the volume determined by the previously listed ratios. Samples were incubated at room temperature then placed on a magnet stand until the solution was clear. The supernatant was discarded, and freshly made 80% ethanol was added to each sample. This ethanol purification step was performed twice to remove impurities. All ethanol was removed and beads were allowed to dry for one minute before ddH2O was added and the samples were removed from the magnet. Samples incubated for 5 minutes at room temperature, then were placed back on the magnet stand. The eluted ladder was moved to another tube and the beads were discarded. The cleaned samples were visualized using gel electrophoresis on a 2% agarose gel run at 100 V and imaged on UVP GelStudio Plus (Analytik Jena, Jena, Germany). The gel image was edited in GIMP 2.10.34 to improve clarity.

### Experiment 2

We next tested the bead protocol on museum material and optimized the recipe to generate the highest quality extract possible. Salt concentration can affect the coprecipitation of impurities; for this reason, we tested different molarities of NaCl. We combined alterations in NaCl molarity (1 M, 2 M, 2.5 M) with variable PEG-8000 concentrations (20% and 21%) and bead ratios (1.2x and 1.5x), selected based on the results of the previous experiment. We tested each recipe on 24 museum specimens belonging to 6 orders and 12 insect families with collection dates ranging from 1997 to 2014 (S1). Large-bodied specimens were chosen to ensure high DNA yields and allow comparison across treatments. Each specimen underwent a non-destructive cell lysis procedure—whole bodies were submerged in 600 μL of cell lysis buffer made in-house (S2) and Proteinase K at a concentration of 0.1 mg/ml. Samples were lysed for 16 hours at 55°C on a shaker plate set to 80 RPM. 25 μL of lysate was then removed from the samples and transferred to a 96-well PCR plate.

30 μL or 37.5 μL of each bead solution was added to 25 μL of lysate, depending on the bead ratio being tested. Samples were incubated at room temperature for 5 minutes then incubated on an Alpaqua 96S Super Magnet (Alpaqua Engineering, Beverly, MA, USA) for 5 minutes. Once the beads were bound, the supernatant was discarded using a Rainin BenchSmart semi-automated pipetting machine (Mettler Toledo, Greifensee, Switzerland). 150 μL of fresh 80% ethanol was added using the BenchSmart and samples were incubated in ethanol for 1 minute, then ethanol was discarded. This step was repeated one more. The beads were allowed to dry for 1–3 minutes following final ethanol removal. The plate was then removed from the magnet and 27 μL of ddH2O water was added. The plate was lightly vortexed to homogenize the beads and then briefly spun down before being placed in an incubator at 37°C for 5 minutes. The plate was then lightly vortexed once more and quickly spun down, ensuring the beads remained homogenous, and incubated for another 5 minutes at room temperature. The plate was set on the plate magnet until the liquid was clear, generally 3 minutes, and then the eluted DNA was transferred to another plate. 2 μL of DNA elute was quantified using the Nanodrop 2000c spectrophotometer (Thermo Fisher Scientific) to obtain estimates of DNA concentration as well as 260/230 and 260/280 ratios; these ratios were used as an indication of the quality of the extract.

PCR was performed to test the effect of different variables on amplification success. The 658 bp mitochondrial COI gene was targeted using the “universal” Folmer primers (Table 1). A second amplicon—a “mini-barcode” region— was targeted using a modified version of the COI-CFMRb reverse primer (herein called cgCFMRb) from Jusino et al. 2019, which amplifies the first 181 bp of the COI gene. PCR was performed in a multiplex reaction to amplify both amplicons simultaneously. PCR reactions consisted of 0.15 μL of 5 U/μL Invitrogen *Taq* DNA polymerase (Thermo Fisher Scientific), 1.5 μL of Invitrogen 10x PCR buffer (Thermo Fisher Scientific), 0.75 μL of 50 mM Invitrogen MgCl2 (Thermo Fisher Scientific), 0.6 μL 10 mg/ml BSA (Sigma-Aldrich, St. Louis, MO, United States), 0.3 μL of 10 mM dNTP (Promega, Madison, WI, United States), 0.45 μL of both 10 μM LCO1490 and HCO2198, 0.15 μL of μL dgCFMRb and 8.65 μL of ddH2O, for a 15 μL reaction. PCR conditions were as follows: 94°C for 1 minute, followed by 35 cycles of 1 minute at 94°C, 1:30 minutes at 50°C and 1 minute at 72°C. This was followed by a final extension phase at 72°C for 7 minutes. The reaction was performed on a BioRad T100 thermocycler (BioRad Laboratories, Hercules, CA, United States). Gel electrophoresis was used for visualization and run on a 2% agarose gel at 100 V then imaged using the UVP GelStudio Plus. Gel images were edited in GIMP 2.10.34 to improve clarity and contrast. Amplification success was determined by the presence or absence of bands of the corresponding size on the gel (S1).

**Table 1.**
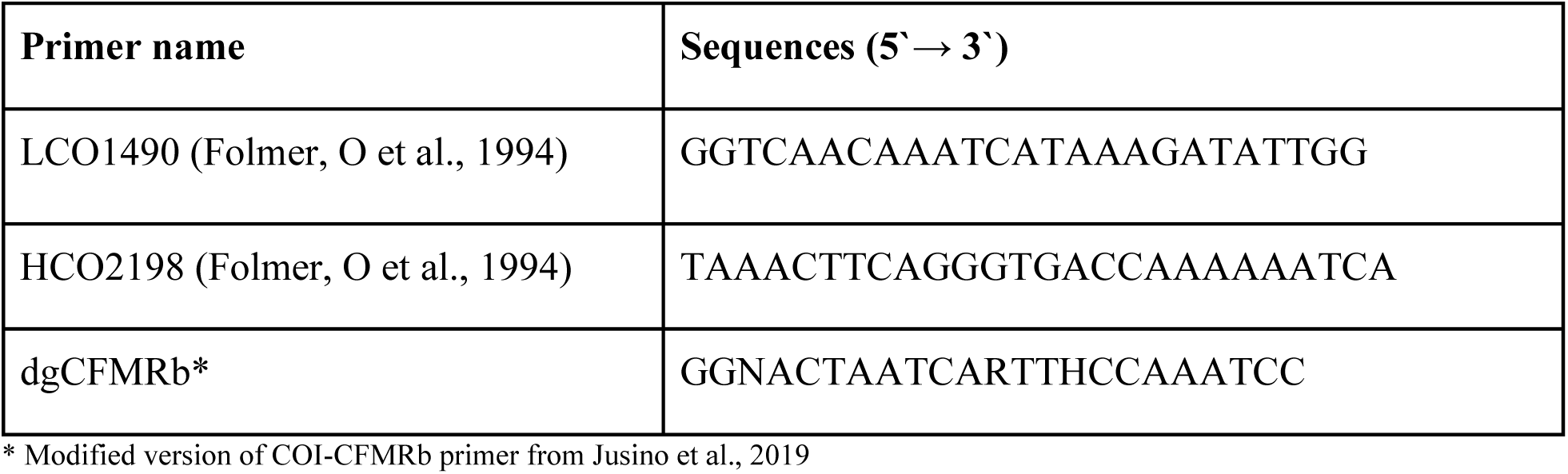
Primer sequences used to amplify regions of COI.

**Table 2.** Summary of results for P. andrewsi samples across methods. DNA yield, 260/230 and 260/280 are median values. Amplification success is the number of bands visible following gel electrophoresis. Qiagen DNeasy resulted in the highest quality extractions. In terms of amplification success, the bead-based method was comparable despite being of slightly lower quality.

|  | DNA yield | 260/230 | 260/280 | PCR success |
| --- | --- | --- | --- | --- |
| Qiagen DNeasy | 16.70 ng/μL | 2.18 | 1.86 | 5 |
| Qiagen Puregene | 4.54 ng/μL | 1.66 | 1.91 | 4 |
| Bead-based method | 9.34 ng/μL | 1.62 | 1.96 | 5 |
| HotShot | 5.02 ng/μL | 1.02 | 2.24 | 1 |

Five of the 24 samples resulted in low DNA yields across all treatments, with median values less than 20 ng/μL. This results in unreliable 260/280 and 260/230 ratios. Because the yield was low across all treatments, and therefore not a result of variability in bead recipes, the samples were removed for analyses. To account for the remaining variability that existed between samples, we employed mixed-effects modeling to assess the importance of NaCl molarity, PEG-8000 concentration, SPRI bead ratios, and specimen age on DNA yield, purity, and amplification success. DNA concentration, 260/280 values, and 260/230 values were log-transformed prior to analysis. The collection year was also centered to improve normality. Model fitting was performed using maximum likelihood with the lme4 package (Bates et al. 2015). Linear mixed-effects models were fitted for DNA concentration, 260/280, and 260/230 individually using function *lmer*. Logistic regression was performed for amplification success using generalized mixed models with a logit link using function *glmer*.

Null models were constructed for each response using sample ID as a random intercept. We then built individual models for each fixed effect and iteratively added additional fixed effects to the best-performing individual model. This process continued until no significant improvement was observed. Models were compared using the Akaike Information Criterion (AIC) and Bayesian Information Criterion (BIC). Likelihood ratio tests (LRT) were used to test model improvements against the null model. Diagnostic plots were examined using the package DHARMa, which produces scaled, interpretable residuals for complex models using simulation (Hartig 2017).

Functions *simulateResiduals*, *plotQQunif* and *plotResiduals* were used for this purpose. A receiver operating characteristic (ROC) curve was constructed for the logistic regression model to assess model performance using function *roc* from the package pROC (Robin et al. 2011).

### Experiment 3

We tested our optimized approach against more commonly used extraction methods to ensure effectiveness. As previously discussed, the most widely used method is the Qiagen DNeasy kit; while the cost is prohibitive for large-scale projects, this kit produces high-quality DNA and served as the “best-case” extraction. Next, we performed a salt precipitation method using the Qiagen Puregene kit. These approaches are inexpensive but require multiple steps that can be lengthy as well as lead to additional shearing. The third method we tested was the HotShot extraction approach (Truett et al. 2000), a rapid and low-cost alkaline extraction approach that does not include a DNA purification step. HotShot has recently been adopted in high-throughput barcoding studies targeting Diptera (Srivathsan et al. 2021). These methods, along with our bead-based extraction protocol, were used on leg tissue from museum specimens - the beetle species *Pseudocotalpa andrewsi* and the fly species *Hybomitra sonomensis,* with five replicates within each species for each approach. All specimens were collected between 1977 and 1986.

Each method began with the mastication of leg tissue using a pestle. Qiagen DNeasy was performed according to the manufacturer’s instructions. Salt precipitation was conducted using the Qiagen Puregene tissue extraction kit according to the manufacturer’s instructions. HotShot was performed based on the protocol in Truett et al., 2000 - samples were heated at 95°C for 30 minutes in a highly alkaline cell lysis buffer consisting of 25 mM NaOH and 0.2 mM EDTA. After incubation, samples were cooled and a neutralization buffer consisting of 40 mM of Tris-HCl was added to bring the pH to a neutral range. No further DNA purification steps were involved in the protocol. Bead-based extractions were performed using a solution consisting of 21% PEG-8000, 1 M salt, and 1.5x bead ratio. DNA was quantified using a Qubit 2.0 fluorometer (Thermo Fisher Scientific), quality was assessed using a Nanodrop 2000c spectrophotometer, and fragment sizes were assessed with a Tapestation system (Agilent Technologies, Santa Clara, CA, USA).

To further assess quality, we performed PCR targeting a mini-barcode. *H. sonomensis* samples were amplified using LCO1490 and dgCFMRb as in Experiment #2. Uni-MinibarF1 (5’-GAAAATCATAATGAAGGCATGAGC-3’)(Meusnier et al. 2008) and Uni-MinibarR1 (5’-TCCACTAATCACAARGATATTGGTAC-3’)(Meusnier et al. 2008) primers were used to amplify *P. andrewsi* samples as LCO1490 can perform poorly in order Coleoptera. PCR reactions consisted of 0.15 μL of 5 U/μL Invitrogen *Taq* DNA polymerase (Thermo Fisher Scientific), 1.5 μL of Invitrogen 10x PCR buffer (Thermo Fisher Scientific), 0.75 μL of 50 mM Invitrogen MgCl2 (Thermo Fisher Scientific), 0.6 μL 10 mg/ml BSA (Sigma-Aldrich), 0.3 μL of 10 mM dNTP (Promega), 0.3 μL of 10mM forward and reverse primers and 8.65 μL of ddH2O. PCR thermocycler conditions were the same as in Experiment #2. Amplification success was assessed using gel electrophoresis with 2% agarose run at 100V.

ANOVA was performed to test the effect of the extraction method on DNA concentration for *H. sonomensis* and *P. andrewsi* samples separately. Tukey’s honest significance test (Tukey’s HSD) was performed to assess the significance of concentration differences between extraction methods. Further analyses were performed exclusively on *P. andrewsi* data due to the extremely low DNA concentrations from *H. sonomensis* samples. ANOVA and Tukey’s HSD were used to test differences in extract quality, based on 260/280 and 260/230 ratios. Amplification success was determined based on gel electrophoresis results. TapeStation curves were edited using InkScape 1.3.2.

### Experiment 4

The previous three experiments led us to a bead-based protocol that paired a gentle cell lysis step with nucleic acid purification using a 1.5x ratio of beads prepared in a solution of 1 M NaCl and 21% PEG. We tested the applicability of the approach on large-scale projects with different types of samples. Samples were provided by the California Academy of Sciences, the California Department of Food and Agriculture, the University of California Riverside, and an independent collaborator. All samples were plated in Qiagen 1.2 mL collection microtubes in 96-well plates and consisted of either leg tissue or whole bodies. Samples that were non-destructively extracted by soaking in cell lysis buffer were plated into separate categories to allow different treatments. Specifically, samples were split into three size categories (small, medium, and large) and were split by degree of sclerotization. These categories allowed us to vary the volume of cell lysis buffer, Proteinase K concentration, and elution volume as well as lysis time (S2).

Plates that contained tissue required homogenization before lysis. This was done by adding two 3.2 mm stainless steel beads to each well along with a standard volume of carbide sharp particles (BioSpec, Bartlesville OK, USA) measured using a stainless steel 1/32^nd^ teaspoon measure. We designed a plate insert that can be manufactured using 3D printing that allowed easier addition of beads and particles to each well (S3, S4). This was sterilized between each plate. Cell lysis buffer and Proteinase K were added following the addition of homogenization materials and tissue was homogenized using a Tissuelyzer II (Qiagen). For plates containing whole specimens, cell lysis buffer and Proteinase K was added directly to each well. Plates were then incubated at 55°C, with incubation time varied by the type of material (S2).

Lysate was transferred from deep well plates to PCR plates using 200 μL filtered tips. 25–50 μL of lysate, volume dependent on yield expectations, was transferred into a 96-well PCR plate containing the SPRI bead solution. For specimens lysed non-destructively, samples were immediately rinsed then stored in 95% ethanol to be returned to collections. Purification using SPRI beads occurred as described in previous sections. A detailed protocol can be found in S2. Following purification, 12 wells across each plate were quantified using the Nanodrop 2000c spectrophotometer to assess general extraction success. Negative controls were additionally quantified.

To assess the overall success of extraction, random wells on each plate were quantified using Nanodrop 2000c. ANOVA was performed on the log-transformed DNA concentration data to test the difference between tissue types - either subsampled leg tissue or non-destructive whole. Linear models were constructed separately for both extraction methods to assess the relationship between sample age and yield.

## Results

### Experiment 1

In typical bead protocols used for size selection, lower bead ratios target large DNA fragments and result in the removal of small DNA fragments. Our goal was to alter this behavior to obtain small fragments even when using low bead ratios, reducing the cost by reducing the volume of beads necessary. We found that fragment retention increased with the percentage of PEG. At 18% PEG, 100 bp fragments were faint; paired with a 1.4x ratio and below, the 100 bp fragments were lost. At 19% PEG, the 100 bp fragments were lost when the ratio was below 1.2x. At 20% PEG, the 100 bp fragments were lost when the ratio was below 1.0x. Both 20% and 21% PEG concentrations retained 100 bp fragments up to 0.8x ratio, although the bands are faint. 22% looked similar to 21% in fragment size retention (Fig. 1).

**Figure 1.**
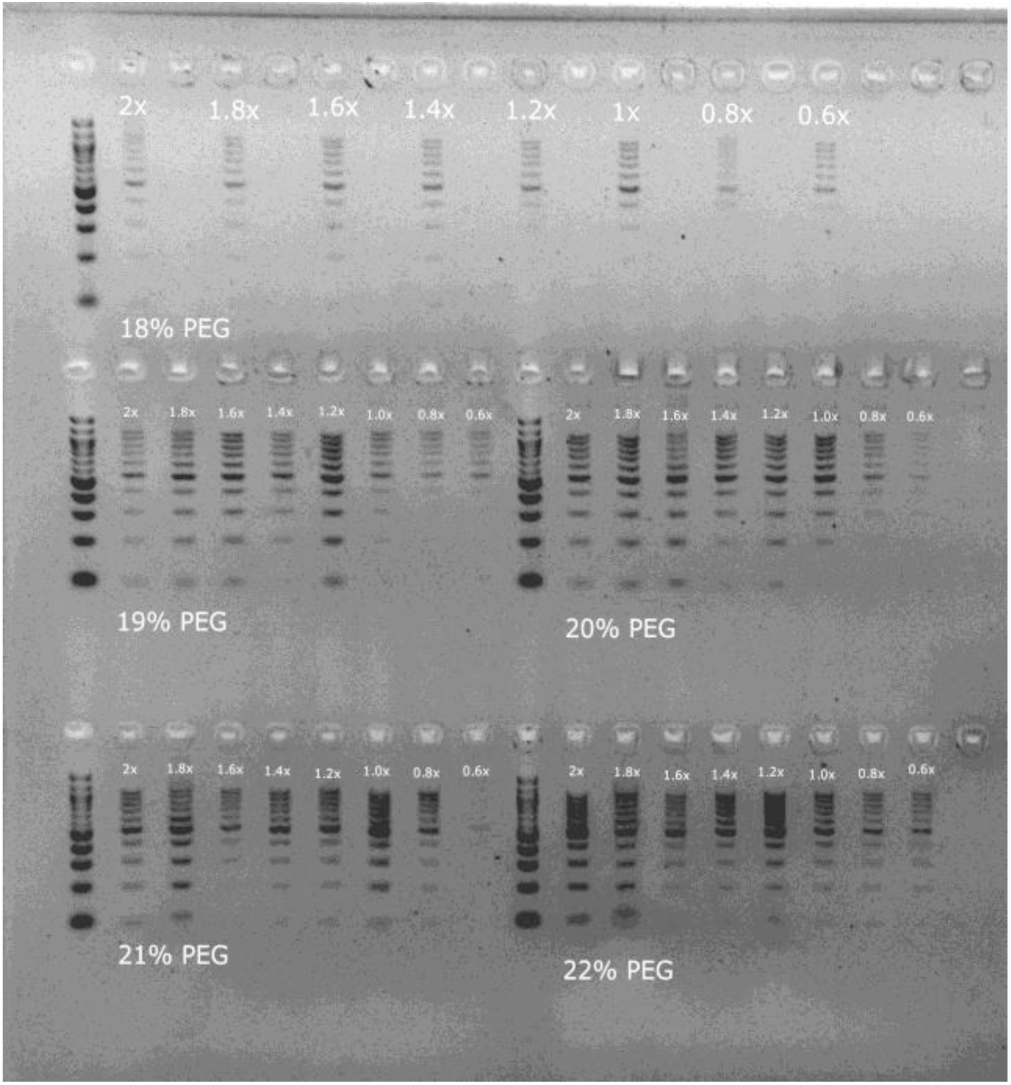
Effect of PEG-8000 concentration on size selection. Gel image depicting the outcomes of bead purification performed on a diluted GeneRuler ladder sample. The figure is organized into blocks, each representing a different PEG-8000 concentration ranging from 18% to 22%. Within each block, columns correspond to varying ratios of beads to sample volume. The results demonstrate that, as the PEG-8000 concentration increases, smaller DNA fragments are more likely to be retained.

Because high PEG ratios can result in increased co-precipitation of impurities, we opted for 21% PEG, which would allow us to reduce our bead ratio while retaining small DNA fragments.

### Experiment 2

Five samples were removed from further analyses due to failed extractions across all treatment types. Following the removal of these samples, the median DNA concentration across samples and treatments was 149.5 ng/μL but varied widely by sample (S1). The lowest median yield was 143.15 ng/μL and was produced by a solution of 20% PEG and 1M NaCl using a 1.2x bead ratio. The highest median yield was 213.25 ng/μL, produced by a solution of 21% PEG and 2 M NaCl using a 1.5x bead ratio (S1).

Using a mixed-effects model, we found that bead ratio, PEG concentration, and NaCl molarity were not significant predictors of DNA concentration. The year in which the sample was collected had the strongest effect on yield (β = 0.441, t-value = 2.02). The best-fitting model included an interaction between year and NaCl (χ^2^ = 31.688, p-value = 6.85e-06; S1) with a 2M NaCl solution having the largest effect (Fig. 2). However, diagnostic checks revealed significant deviations in quantile-quantile (QQ) residuals and residuals vs. predicted plots, indicating potential violations of model assumptions (S1). These deviations suggest that the results should be interpreted with caution.

**Figure 2.**
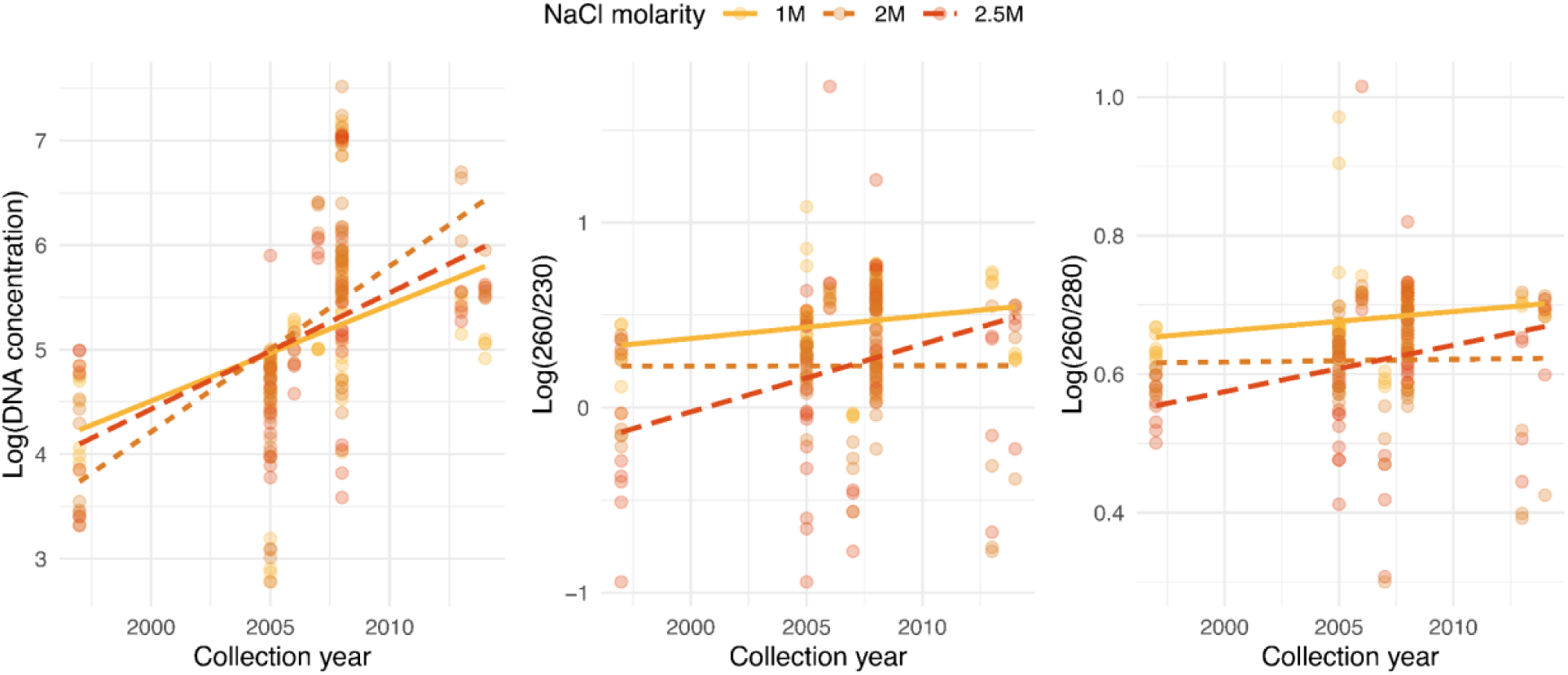
Interaction between NaCl molarity and collection year in prediction of DNA concentration, 260/230 and 260/280 ratios. Year had a strong linear relationship with DNA concentration. Younger samples resulted in increased DNA yields, with 2 M NaCl producing the steepest slope. NaCl molarity had the greatest effect on extract quality, with 2 M and 2.5 M NaCl yielding the lowest quality extracts. 2.5 M had a strong interaction with sample age, with a steep decrease in quality in older samples.

NaCl molarity was the most influential factor in 260/280 and 260/230 values with higher molarities negatively affecting both ratios. All models including NaCl as a feature significantly improved model fit compared to the null model (S1). 1 M NaCl produced the highest values for both 260/280 and 260/230, with a median of 260/280 of 1.97 and a median 260/230 of 1.62 (S1). Molarity differences had the most pronounced effect on 260/230 ratios, with a median of 1.33 for 2M solutions and 1.36 for 2.5M solutions (2M: β = -0.21214, t-value = -4.583; 2.5M: β = - 0.26777, t-value = -5.784). Inclusion of interaction between year and NaCl produced the best-fitting model for 260/280 (χ^2^ = 44.352, p-value = 1.96e-08; S1) and 260/230 ratios (χ^2^ = 53.191, p-value = 3.07e-10; S1); this was largely driven by 2.5M solutions in which the purity of extracts improved significantly as collection year became more recent (Fig. 2). Once again, the inclusion of year caused significant residual deviations in the model predicting 260/280, indicating potential model misspecification. A model including NaCl as the only feature showed no major assumption violations and still provided a significantly better fit than the null model. No clear model violations were observed when including NaCl and year as fixed effects in the prediction of 260/230 values. NaCl can be most confidently identified as influential in the differences in purity across models while the influence of age is unclear.

Extractions using the 1 M bead solution produced the highest number of full and mini-barcodes. Extractions using a 1.2x bead ratio with a bead solution containing 1 M NaCl and 20% PEG produced the highest number (n = 10) of full barcodes. All other extractions using 1 M solutions produced 8 full barcodes, while the other treatments at higher NaCl molarities produced fewer (Fig. 3). Models failed to converge when using amplification success of the full COI region due to overall low success. For the 181 bp mini-barcode region, we found that a model including NaCl better predicted amplification success than the null model (χ^2^ = 10.250, p-value = 0.006).

**Figure 3.**
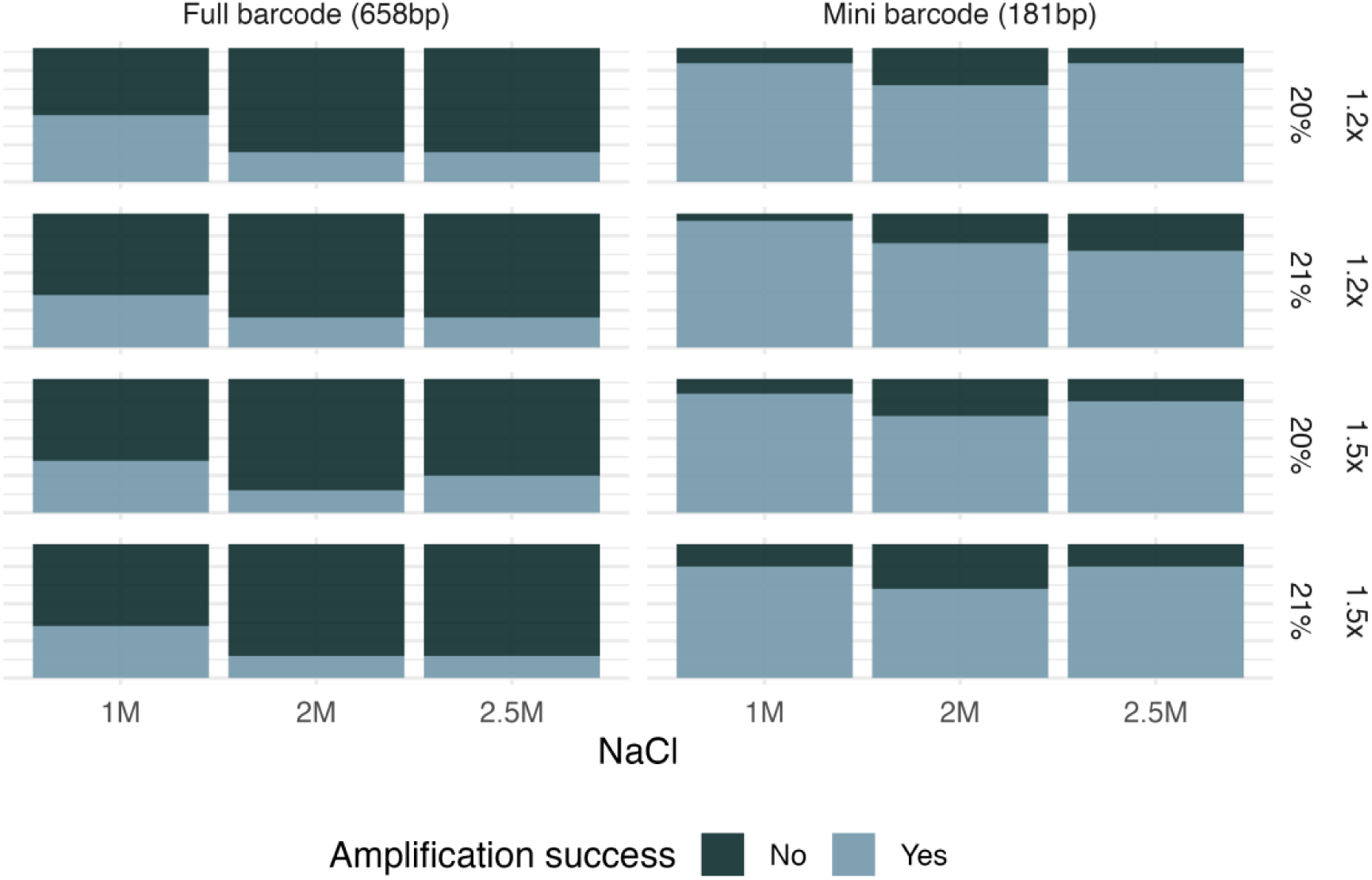
Amplification success of the 658 bp full COI gene and the 181 bp mini-barcode using different bead ratios, PEG-8000 concentrations, and NaCl molarities. Color indicates amplification success, with green representing failed amplification and blue representing successful amplification. Each row shows results from the different combinations of PEG-8000 concentrations (20% and 21%) and bead ratios (1.2x and 1.5x). Columns show different molarities of NaCl, grouped by targeted amplicon. A combination of 20% PEG and 1 M NaCl at a 1.2x bead-to-lysate ratio produced the highest amplification success for the full barcode and all 1 M solutions amplified better than other molarities regardless of PEG or bead ratio used.

Neither ratio nor PEG concentration was significant in predicting amplification success in comparison to the null model.

### Experiment 3

The Qiagen DNeasy kit resulted in the highest DNA concentrations with a median concentration of 16.70 ng/μL for beetles and 1.64 ng/μL for fly samples (Fig. 4). The bead-based method had the second highest median concentration for beetles, at 9.34 ng/μL. Fly samples resulted in much lower concentrations for other methods and these concentrations differed significantly when compared to DNeasy yields (Fig. 4).

**Figure 4.**
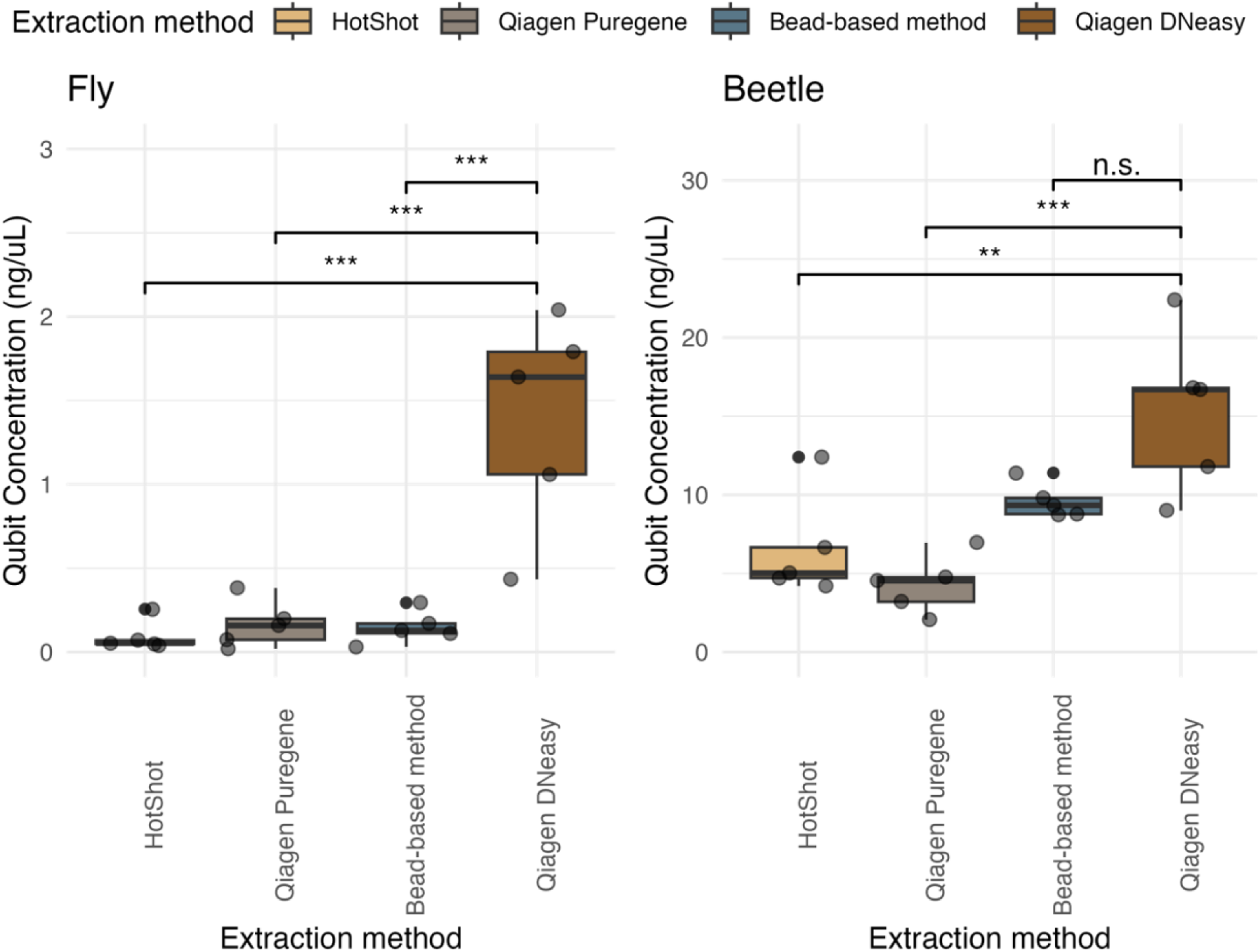
Comparison of DNA concentrations produced from the fly samples (H. sonomensis) and beetle samples (P. andrewsi) using four different extraction methods. Extraction approaches included Qiagen DNeasy, Qiagen Puregene, HotShot, and the bead-based method outlined in this manuscript. Each method is represented by a different color. Qiagen DNeasy resulted in the highest concentration and was significantly different from all methods when extracting H. sonomensis samples. Qiagen DNeasy was significantly different from HotShot and Qiagen Puregene when extracting P. andrewsi samples, but not significantly different from the bead-based method.

260/280 and 260/230 ratios were not reliable for *H. sonomensis* samples because of the low DNA concentration, so ratios were compared only for beetle samples. There were significant differences between extraction methods and 260/280 ratios (F-statistic = 5.649, p-value = 0.008), with the bead-based approach producing the highest values. The difference between 230/260 values across extraction methods was more defined (F-statistic = 19.55, p-value = 1.34e-05).

Results differed significantly among all methods except between the bead-based method and the Qiagen Puregene kit. The difference between HotShot and DNeasy was most pronounced, with the mean 260/230 ratio for HotShot equaling 1.02 compared to DNeasy at 2.18 (Estimate = 1.19, p-value = 5.96e-06).

Low concentrations also produced unreliable Tapestation results for *H. sonomensis*, so these samples were not included. For *P. andrewsi*, the DNeasy kit and the bead-based extraction protocol resulted in the largest fragment sizes; the largest mean fragment size for DNeasy was 452 bp followed by the bead-based approach at 403 bp (Fig. 5). The modified Puregene protocol followed with a 363bp peak and HotShot resulted in the shortest fragments, with a mean of 282 bp.

**Figure 5.**
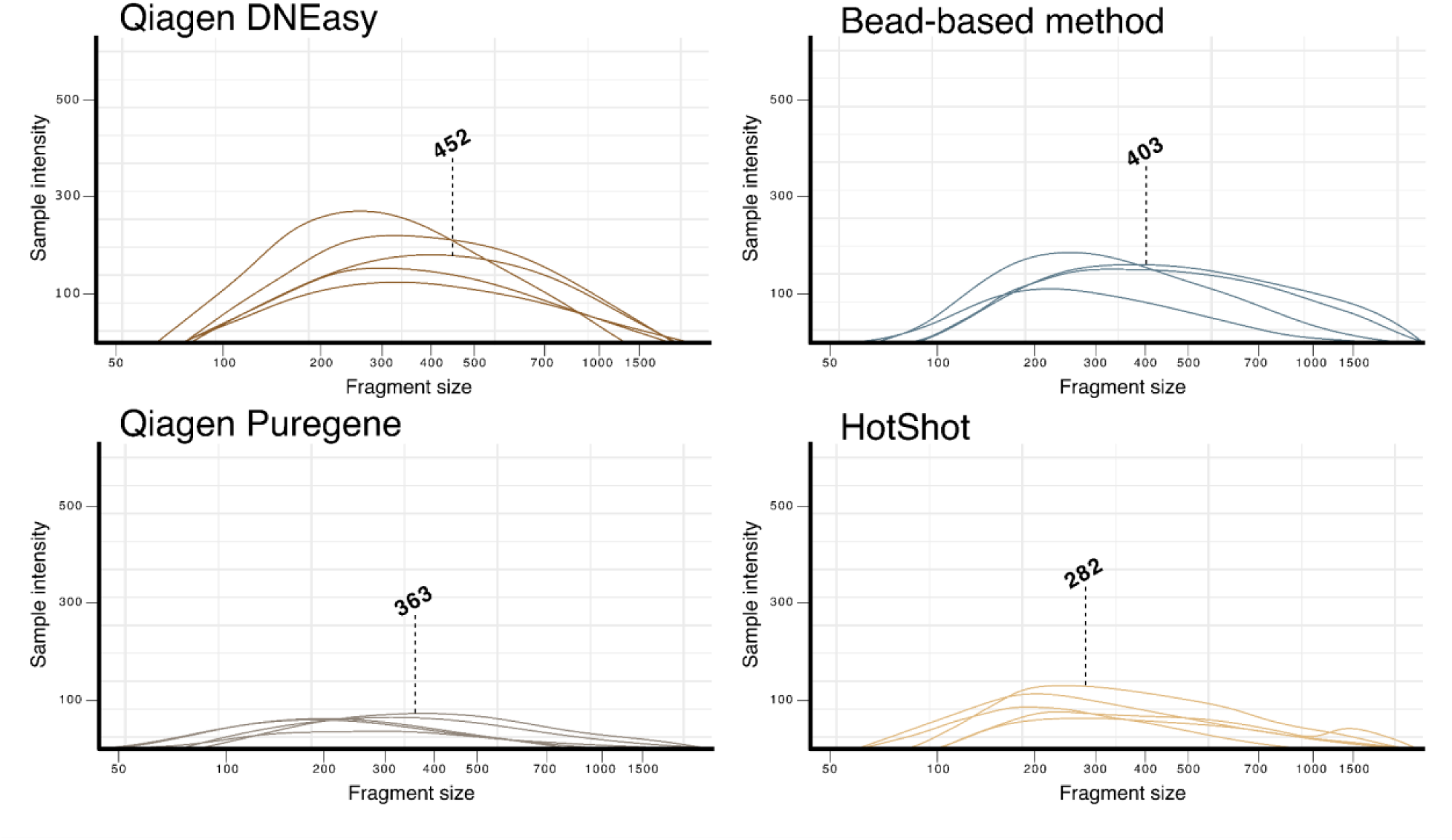
Tapestation curves for the four extraction methods tested on the beetle samples (P. andrewsi). The dashed line and number show the largest mean fragment size for each approach. Qiagen DNeasy retained the highest concentration of large fragments, followed closely by the bead-based method. The HotShot approach produced the smallest fragments.

Qiagen DNeasy produced five bands for *P. andrewsi* and three bands for *H. sonomensis*. The bead-based method was slightly more successful, producing five bands for *P. andrewsi* and four bands for *H. sonomensis*. Qiagen Puregene worked well for *P. andrewsi* samples, amplifying four samples, but was largely unsuccessful for *H. sonomensis* samples, producing only one band. HotShot was the least successful method, with one band for *P. andrewsi* samples and two bands for *H. sonomensis* samples.

### Experiment 4

We extracted DNA from 3,786 specimens belonging to 11 different orders. The samples were collected from 1972 to 2024 and had a mean age of 19 years old. We found that altering the lysis time, Proteinase K concentration, and reagent volumes based on tissue types greatly benefited extraction yield, and we established a standard protocol for this (S2). By using a BenchSmart, nucleic acid purification using SPRI beads took roughly 45 minutes following cell lysis. Including the time needed to transfer lysate, the protocol takes approximately one hour in total. Excluding lab consumables, the extraction approach costs 10.7 cents in cases where 50 μL of lysate was used, and 6.5 cents in cases where 25 μL of lysate was used. This was using a 1.5x ratio bead clean; we found that a 1.2x clean was often suitable but opted for a higher ratio out of caution. Using a 1.2x bead ratio would further reduce costs to 9.1 and 5.7 cents respectively and concentrate larger fragments but risks loss of small fragments.

Plates were spot-checked using Nanodrop to assess success; the data analyzed was based on 495 samples across all plates (40 96-well plates). Non-destructive methods resulted in the highest yields, with a mean DNA concentration of 52.37 ng/μL and median of 21.4 ng/μL, while homogenized leg tissue resulted in a mean DNA concentration of 12.7 ng/μL and median of 8.1 ng/μL (F-statistic = 74.22, p-value = <2e-16, Fig. 6). There was no linear relationship found between sample age and DNA concentration for either method (S1).

**Figure 6.**
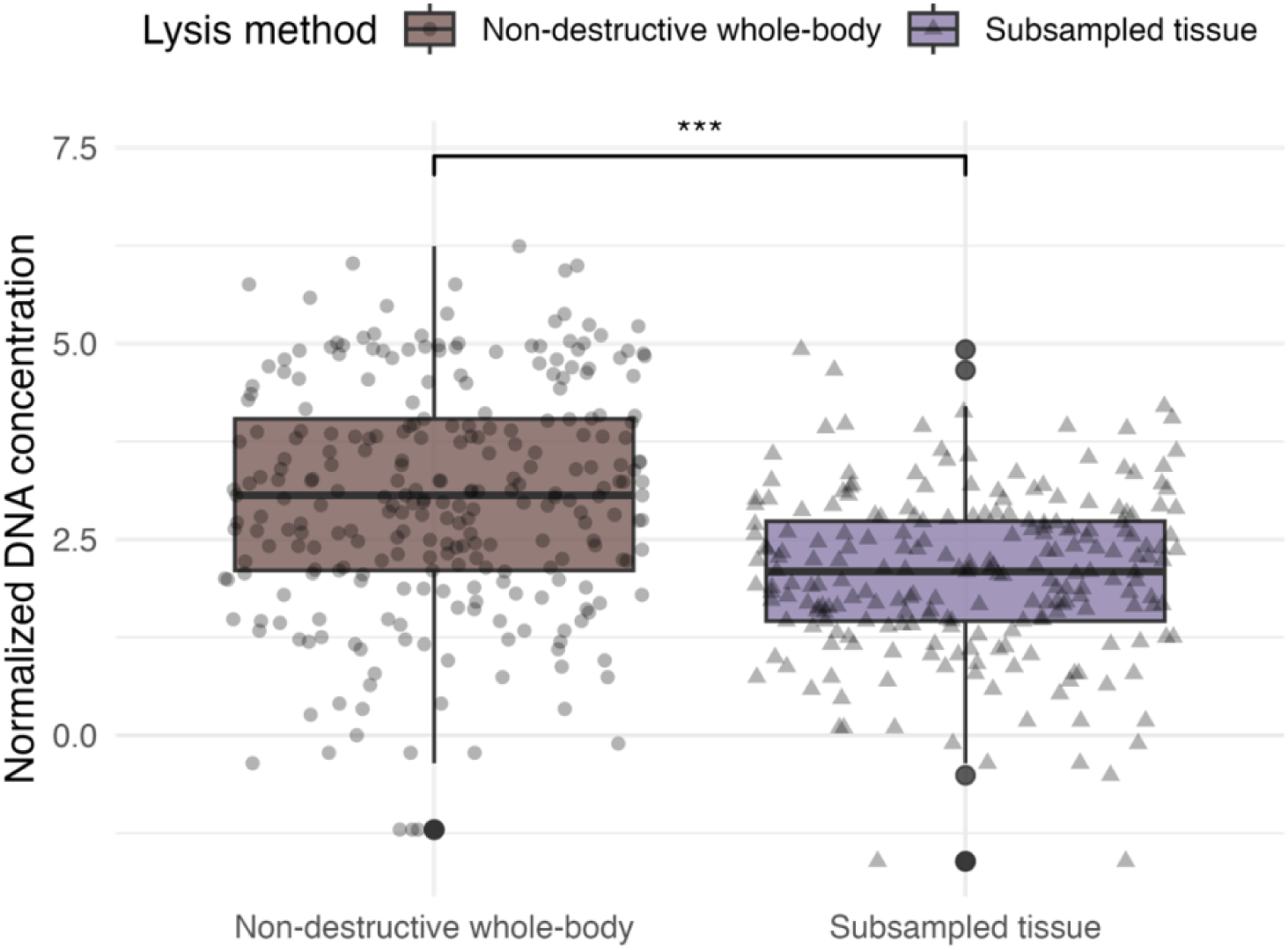
Boxplot showing the impact of input tissue type on normalized DNA concentration. Point shape and box colors represent methods. Non-destructive methods in which whole bodies were soaked in cell lysis buffer resulted in the highest median and mean yields when compared to extracts using subsampled leg tissue (p-value = <2e-16).

## Discussion

Our study aimed to test and optimize a cost-effective extraction method to obtain archival-quality DNA from museum specimens across taxonomic groups and tissue types in a high-throughput manner. We found a SPRI bead-based extraction protocol effective in balancing cost and throughput capacity for a wide range of specimens. The approach provides flexibility in the equipment needed and in the reagent volumes, allowing for adaptation to project specifics. Our approach has the potential to greatly improve the ability to obtain DNA from diverse collections, moving us toward the rapid generation of genomic data from museum collections.

### Bead solution optimization

Large-scale museomics projects have partially been hindered by costs. Therefore, a major aim of our study was to minimize extraction costs while maximizing extraction quality. The most expensive component of bead-based approaches is the beads themselves. Prepared bead solutions, such as AMPure XP beads (Danaher Corporation), are expensive and not feasible for large projects. However, it is possible to make custom bead solutions in-house (Rohland and Reich 2012), greatly reducing cost.

When preparing stock beads, polyethylene glycol (PEG-8000) and salt (NaCl) are the two components of the solution most influential in the success of DNA purification and the resulting size selection. PEG primarily serves as a crowding reagent, forcing nucleic acids out of solution and resulting in size selection for larger-sized fragments (Paithankar and Prasad 1991). High PEG concentrations alone have been commonly used in DNA precipitation, especially for plant material (Youssef et al. 2015). A major benefit of using PEG is the reduction in carryover of polysaccharides that can result from alcohol precipitation approaches (Footitt, Awan, and Finch-Savage 2018; Huang et al. 2018). Polysaccharide contamination is a widely documented problem for extracts from plant tissues but is also a major problem for arthropods, as exoskeleton chitin is one of the most abundant naturally occurring polysaccharides (Tharanathan and Kittur 2003).

Because PEG could alter the retention of DNA fragments, and therefore the bead ratio necessary, we began by testing the effect of PEG concentrations. Altering the percentage of PEG in solution directly related to fragment size selection. At lower percentages, small fragments of the DNA ladder were not retained. This is beneficial for the classic use of beads in size selection but, when working with degraded DNA, loss of small fragments is problematic. We found that increased concentrations of PEG retained smaller fragments when using low ratios of beads. This is essential for cost reduction - the less beads necessary, the cheaper the extraction protocol.

PEG causes DNA precipitation in combination with salts such as NaCl which increase the ionic strength of the solution. Salt concentration can be altered to not only improve nucleic acid binding and improve total yield but also to reduce impurities. The NaCl molarity used in bead solutions varies in the literature from 1 M to 2.5 M (Liu et al. 2023, 292; Rohland and Reich 2012; Hawkins et al. 1994) but there is no defined rationale behind the selection of specific molarities. We found that NaCl was the most important factor associated with extract purity, with lower molarities resulting in higher-quality extracts. Across normalized 260/230 and 260/280 values, 1 M NaCl concentration resulted in the best values for each ratio. A 1.2x ratio combined with 20% PEG was best for reducing contaminants. This solution produced the highest number of full COI barcodes, likely due to the purity of the extract and the increased density of larger fragment sizes achieved through bead size selection. However, this combination produced the lowest concentration of DNA, again likely due to size selection, where small DNA fragments would be removed. This could be problematic for older samples where the expectation is that DNA will be highly fragmented. The oldest sample in this experiment was collected in 1997, making it relatively young for a museum specimen. To ensure small fragments are retained, we proceeded with a solution consisting of 21% PEG-8000 instead of 20%. From here, the bead ratio can be altered based on extraction expectations. For younger samples, where high-molecular-weight DNA is expected, lower bead ratios can be used, while for older samples in which significant shearing has likely occurred, higher bead ratios can be used to ensure small fragments are retained. This allows flexibility depending on the project specifics.

### Comparing to other common extraction methods

Following solution optimization, we compared our approach to other widely used extraction methods. Qiagen DNeasy resulted in the highest yields and ideal Nanodrop values, indicating low protein, salt, and polysaccharide carryover. Studies have successfully used this method to produce genomic information for a variety of preserved arthropod specimens, including pinned beetles (Kanda et al. 2015), microhymenoptera (Guzmán-Larralde et al. 2017), and spiders (Wood et al. 2018). This extraction method and other targeted kits are ideal for those new to molecular approaches, who are processing a small number of samples, and/or have abundant funding at their disposal. However, kits are impractical and cost-prohibitive when attempting to process thousands of samples. A quality sacrifice is expected for cheaper methods but is necessary to move towards higher throughput.

We contrasted our SPRI approach and other methods against DNeasy extractions, treating this kit as a “gold standard” comparison. The 260/280 values were near 2.0 for both the bead-based method and the Puregene protocol. 260/280 ratios between 1.8–2.0 are the accepted range for good purity, while lower values indicate protein carryover. Values closer to 2.0 can indicate RNA presence. The median 1.96 value produced from the bead-based protocol may be due to degraded RNA, which would also bind to carboxyl groups on SPRI beads. In PCR-based approaches, RNA is not of great concern. However, if RNA could interfere with downstream applications, an additional RNAse step following cell lysis would easily resolve this concern.

The 260/230 values for all extraction methods except Qiagen DNeasy were lower than optimal. This is very common for arthropods given their polysaccharide content. The HotShot protocol unsurprisingly produced the worst purity readings due to the lack of a purification step.

The SPRI-bead approach was competitive with the DNeasy kit when it came to the retention and concentration of larger DNA fragments, which likely increased the amplification success of the samples; four of five fly samples amplified and all beetle samples amplified using extracts from the bead-based protocol in comparison to three of five flies and four of five beetles for the DNeasy kit. The extracts from HotShot had the smallest fragment sizes and resulted in the lowest amplification success across methods. The HotShot approach is an alkaline-based extraction method that includes high-temperature lysing and a pH that can increase DNA shearing (Rudbeck and Dissing 1998). As stated previously, there is no purification step, meaning all inhibitors and other components released from cells during lysis remain in the elute. These effects likely explain the small fragment sizes we observed and the low amplification success.

Other studies found this approach unsuitable for specimens older than 12 years (Guzmán-Larralde et al. 2017). Due to its cost and simplicity, the HotShot approach is useful for PCR-based studies using younger tissue where high-quality DNA is not necessary but is inadequate for studies targeting older or poorly preserved specimens or studies aiming to produce high-quality archivable genomic DNA.

### High-throughput potential

We were successful in extracting many different types of tissue using this extraction method. We found that by varying Proteinase K concentration, lysis volume and incubation time, we could obtain DNA from very small tissue amounts and from heavily sclerotized specimens that are challenging to extract. Age was not a driving factor in yields for the larger pool of samples, making this approach attractive for museum specimens of various ages. This is especially true because the method targets larger fragment sizes, resulting in the retention of fragments similar to those obtained by the DNeasy kits, and does not appear to induce further shearing that likely occurs in harsher methods such as HotShot.

By using high-throughput tools such as multichannels, semi-automated pipettes or liquid handling machines, this approach can be used to rapidly process thousands of samples. The short hands-on time needed for the bead purification step allows multiple plates to be processed per day. Depending on the bead ratio used, the reagent cost can be as low as 6 cents. 100,000 museum specimens could be extracted for as little as $6,000 in about one year if four plates are extracted per business day. While the BenchSmart does improve the ease of processing many plates per day, we found that using a repeater pipette and multichannel pipettes was also feasible and did not result in quality differences. Equipment such as multichannels and repeater pipettes are more budget friendly and could be purchased with a small grant, allowing a team with limited resources to set up lab space and perform this extraction approach.

## Conclusions

By providing a cost-effective and scalable solution, the SPRI bead-based extraction method optimized in our study effectively addresses key challenges associated with processing diverse specimens from historical collections, including cost constraints, taxonomic breadth, and the accessibility and feasibility of protocols. The adaptability of the protocol allows for modifications in bead ratios, reagent volumes, incubation durations, and tissue types, enabling its application across various sample types and allowing adaption to project specifics such as age, taxon, and preservation state. This versatility makes it an invaluable tool for unlocking the genetic potential of museum archives. As museomics continues to evolve, the adoption of this method could greatly enhance access to genetic data from collections, thereby supporting biodiversity research, conservation efforts, and ecological studies

## Acknowledgments

This study was supported by funding from the California Institute of Biodiversity. In particular, we would like to thank Daniel Gluesenkamp for the generous support. Many people were instrumental in preparing samples. Thank you to the entomology team at the California Academy of Sciences (CAS), specifically Chris Grinter, Julia Betz, Diana Phan, Kate Richards, Levi Ramer and Sarah Nguyen, for their efforts in preparing samples. Additionally, we received samples the California Department of Food and Agriculture, UC Riverside and an independent collaborator – thank you to Glenn Fine, Severyn Korneyev, Julia Perez, Jacob Jones, John Heraty, Paul Rugman-Jones and Christiane Weirauch for your involvement in sample contributions. We are incredibly grateful for the help in the lab from Cecilia Hodson, Rowan Baginsky, and Kate Richards. Many other departments at CAS provided expertise and support; special thanks to Toshiro Chiang from Engineering who designed the plate insert and to Nick Perez and Molly Michelson from the Visualization Studio for producing a companion extraction video. Thank you to our collaborators in entomology at the University of California, Berkeley – Peter Oboyski, Menglin Wang and Rosemary Gillespie – as well as Lydia Smith, Manager of the Evolutionary Genetics Lab for their input.

## Author contributions (CRediT)

**Anna Holmquist -** Conceptualization, data curation, formal analysis, investigation, methodology, project administration, supervision, validation, writing – original draft, writing – review and editing. **Holly Tavris -** Investigation, methodology, writing – review and editing. **Grace Kim -** Investigation, methodology, writing – review and editing. **Lauren Esposito -** Funding acquisition, project administration, writing – review and editing. **Brian Fisher -** Funding acquisition, project administration, resources, writing – review and editing. **Athena Lam -** Conceptualization, funding acquisition, project administration, resources, supervision, writing – review and editing.

## Data availability statement

Additional information on samples can be found in supplementary materials. All other code and data associated with this manuscript are available on Github at https://github.com/ajholmqu/Holmquist2024_MuseumExtractions

