## Supplemental Materials 1 for "Towards large-scale museomics projects: a cost-effective and high-throughput extraction method for obtaining historical DNA from museum insect specimens"

**S1: Supplementary Tables and Figures**

| **Catalog Number** | **Scientific Name** | **Order** | **Family** | **Genus** | **Date** |
| --- | --- | --- | --- | --- | --- |
| CASENT9132408 | Onocosmoecus sp. | Trichoptera | Limnephilidae | Onocosmoecus | 4/19/05 |
| CASENT9132279 | Kathroperla sp. | Plecoptera | Kathroperlidae | Kathroperla | 7/2/05 |
| CASENT9131468 | Rhyacophila sp. | Trichoptera | Rhyacophilidae | Rhyacophila | 6/12/05 |
| CASENT9132912 | Kathroperla takhoma | Plecoptera | Kathroperlidae | Kathroperla | 3/26/08 |
| CASENT9131826 | Kathroperla takhoma | Plecoptera | Kathroperlidae | Kathroperla | 4/17/14 |
| CASENT9131852 | Hetaerina americana | Odonata | Calopterygidae | Hetaerina | 10/31/97 |
| CASENT9130457 | Calineuria californica | Plecoptera | Perlidae | Calineuria | 7/13/13 |
| CASENT9132337 | Calineuria californica | Plecoptera | Perlidae | Calineuria | 2/17/08 |
| CASENT9130978 | Neohermes filicornis | Megaloptera | Corydalidae | Neohermes | 3/3/08 |
| CASENT9132372 | Drunella grandis | Ephemeroptera | Ephemerellidae | Drunella | 4/19/05 |
| CASENT9131493 | Calineuria californica | Plecoptera | Perlidae | Calineuria | 6/12/05 |
| CASENT9130866 | Skwala sp. | Plecoptera | Perlodidae | Skwala | 2/17/08 |
| CASENT9132674 | Dicosmoecus gilvipes | Trichoptera | Limnephilidae | Dicosmoecus | 5/27/07 |
| CASENT9132950 | Calineuria californica | Plecoptera | Perlidae | Calineuria | 2/17/08 |
| CASENT9131716 | Doroneuria baumanni | Plecoptera | Perlidae | Doroneuria | 2/17/08 |
| CASENT9132329 | Rhyacophila sp. | Trichoptera | Rhyacophilidae | Rhyacophila | 5/13/06 |
| CASENT9132686 | Rhyacophila sp. | Trichoptera | Rhyacophilidae | Rhyacophila | 7/25/97 |
| CASENT9131680 | Calineuria californica | Plecoptera | Perlidae | Calineuria | 9/7/08 |

**Table 1. Sample information for Experiment #2.** Includes Catalog Numbers from the California Academy Science’s entomology collection and associated information.

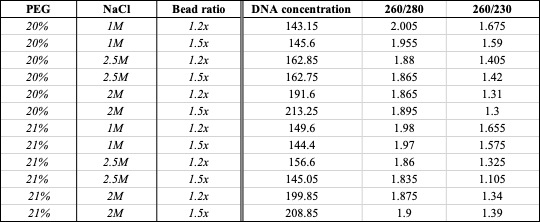

**Table 2. Median DNA concentration, 260/280 ratio and 260/230 values across treatments.** Treatments included a combination of PEG-8000, NaCl and bead ratio. The lowest yield came from 20% PEG, 1 M NaCl and 1.2x bead ratio. This treatment resulted in the highest 260/280 and 260/230 values.

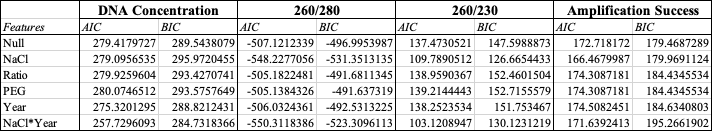

**Table 3. Akaike’s Information Criterion (AIC) and Bayesian Information Criterion (BIC) from all models tested.** Including the NaCl feature consistently improved model fit for purity metrics and amplification success while year was most important for DNA concentration prediction.

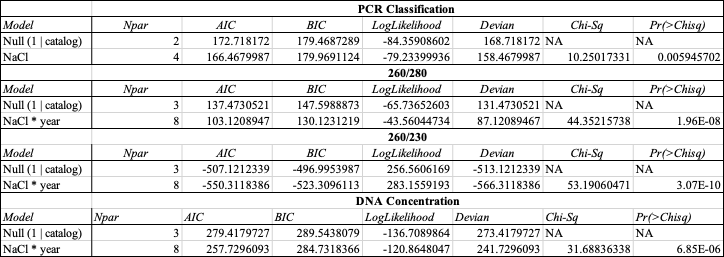

**Table 4. Results from likelihood ratio tests (LRT).** A model including an interaction between year and NaCl most improved model fit for purity metrics and concentration. NaCl alone was the most predictive of amplification success.

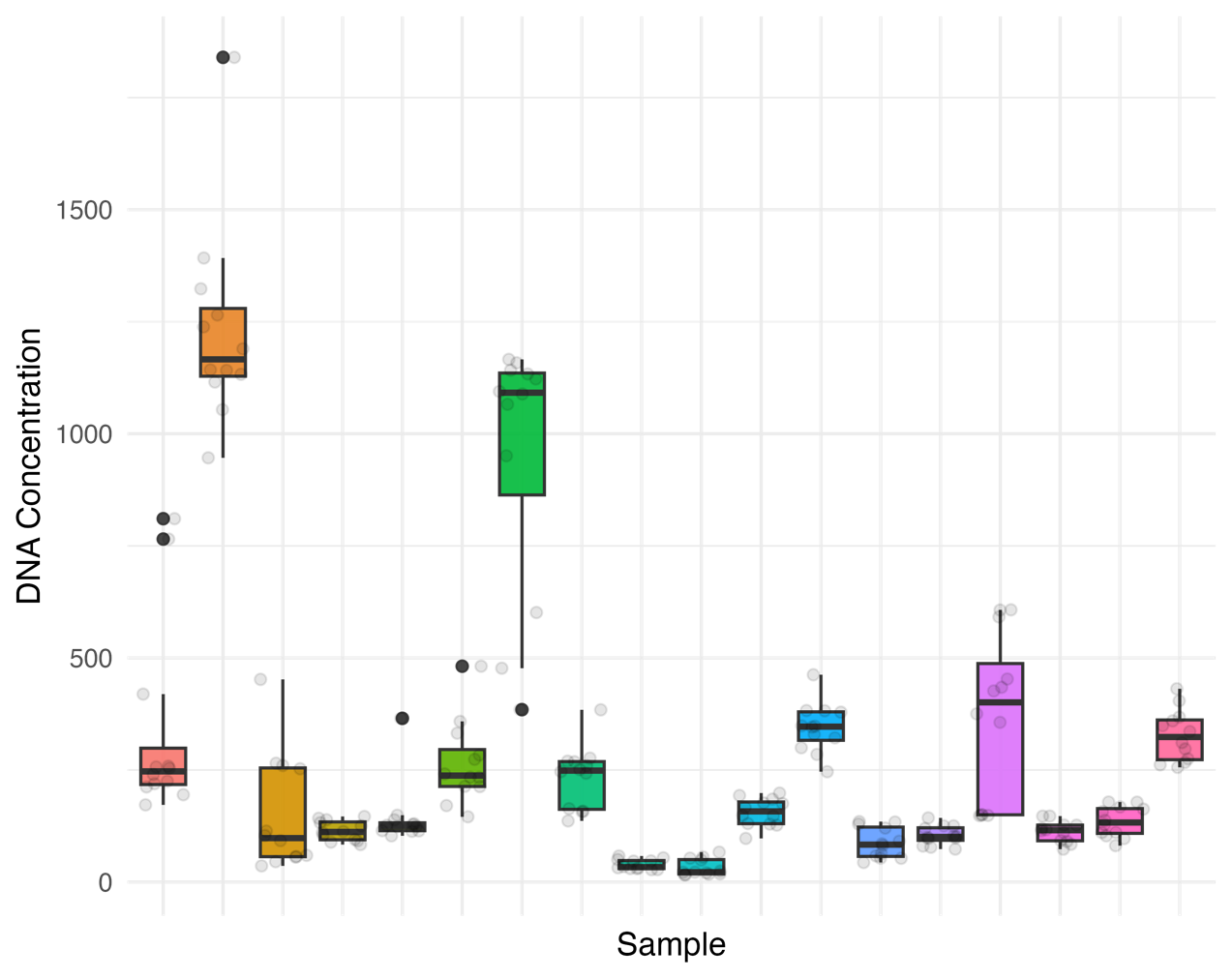

**Figure 1. DNA concentration across samples.** There was high variability in concentration by sample, which led to us choosing mixed-effects models to control for random effects associated with this grouping.

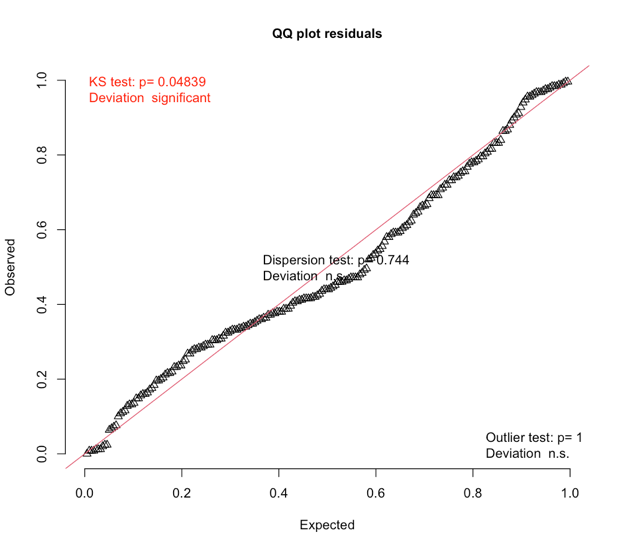

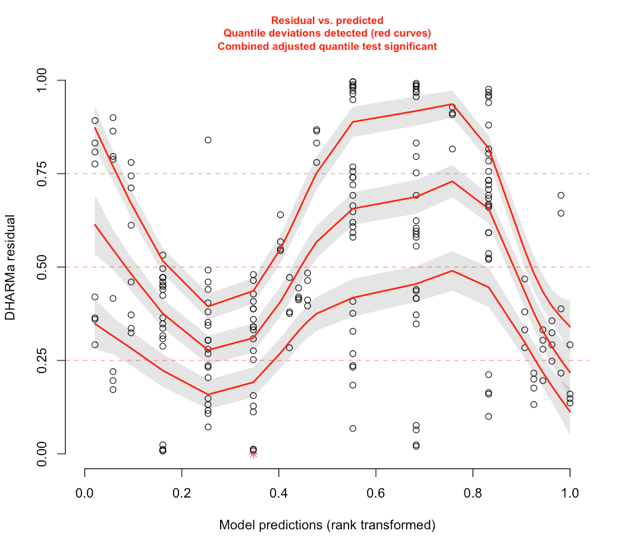

**Figure 2. Model diagnostics including year and NaCl as fixed effects in predicting log-transformed concentration.** The diagnostic plots indicate assumption violations, making the results less reliable.

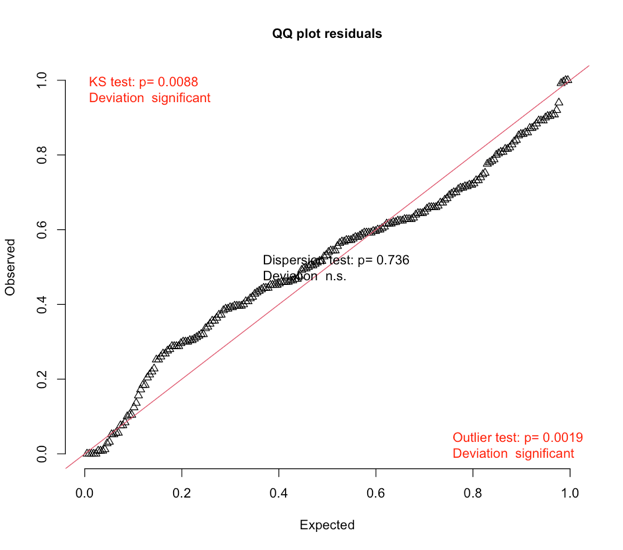

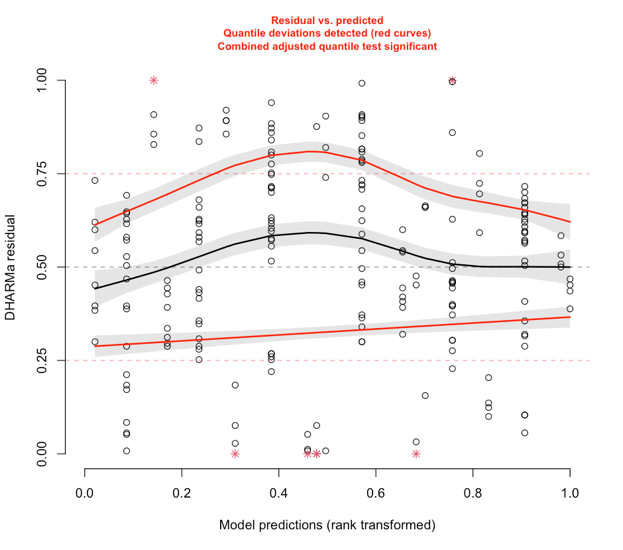

**Figure 3. Model diagnostics for model including NaCl and year as a fixed effect in predicting log-transformed 260/280 ratios.** The diagnostic plots indicate assumption violations, making the results less reliable.

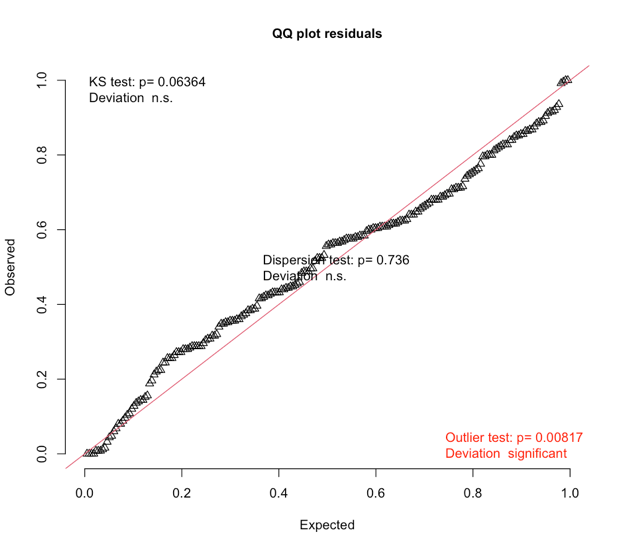

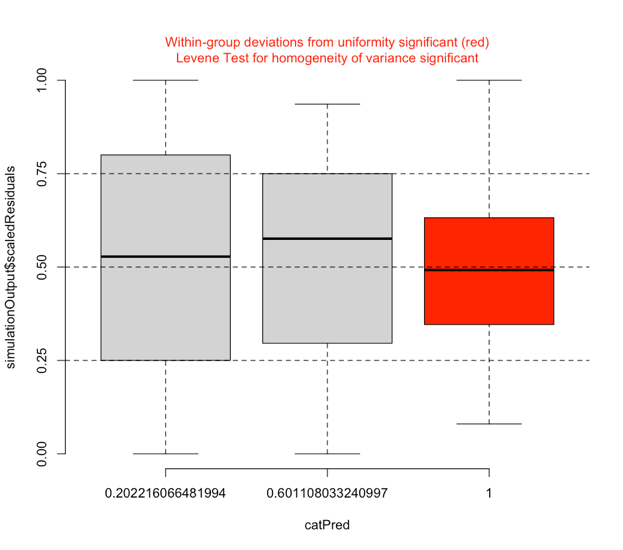

**Figure 4. Model diagnostics for model including NaCl as a fixed effect in predicting log-transformed 260/280 ratios**

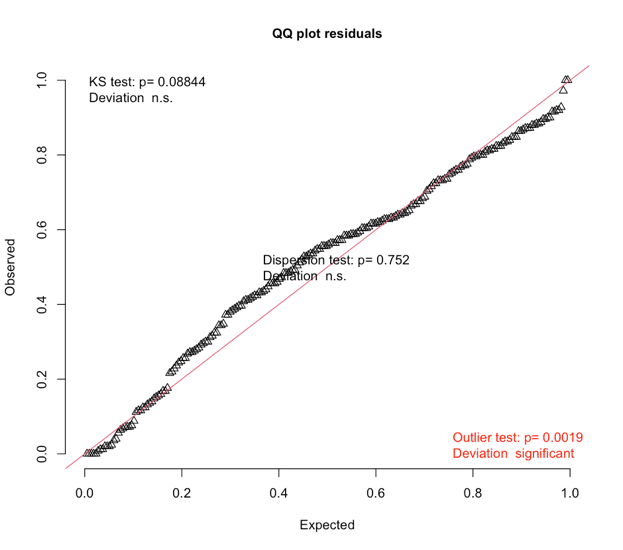

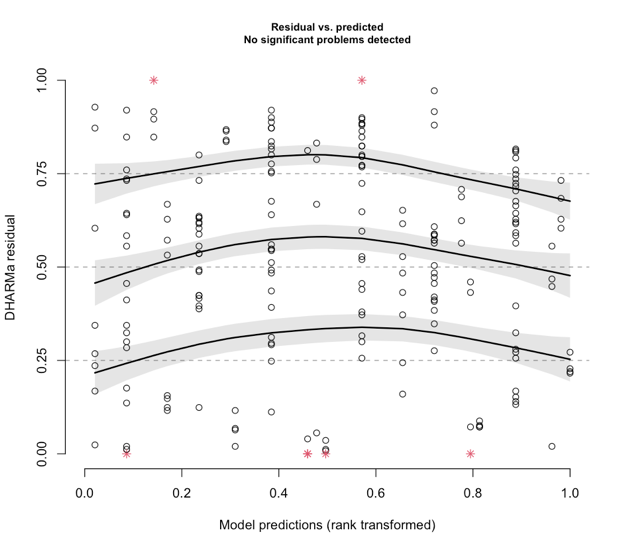

**Figure 5. Model diagnostics for model including NaCl and year as a fixed effect in predicting log-transformed 260/230 ratios.** Diagnostic plots did not indicate assumption violations.

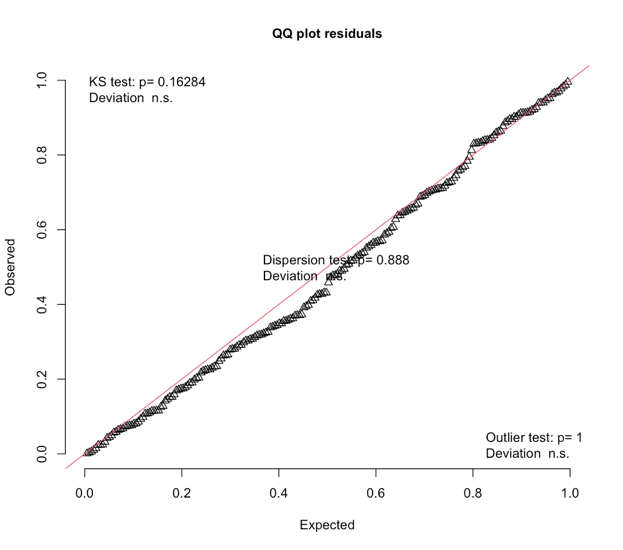

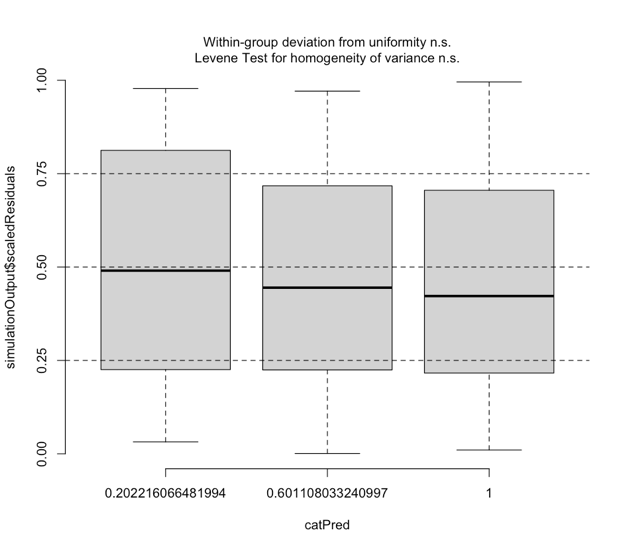

**Figure 6. Model diagnostics for model including NaCl and year as a fixed effect in predicting amplification success for the 180bp mini-barcode**. Diagnostic plots did not indicate assumption violations.

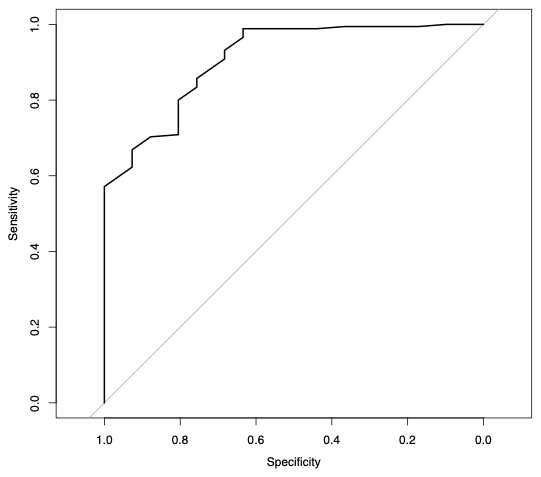

**Figure 7. ROC curve for logistic regression in the classification of PCR success**. Displays relatively high AUC, indicating good model performance.

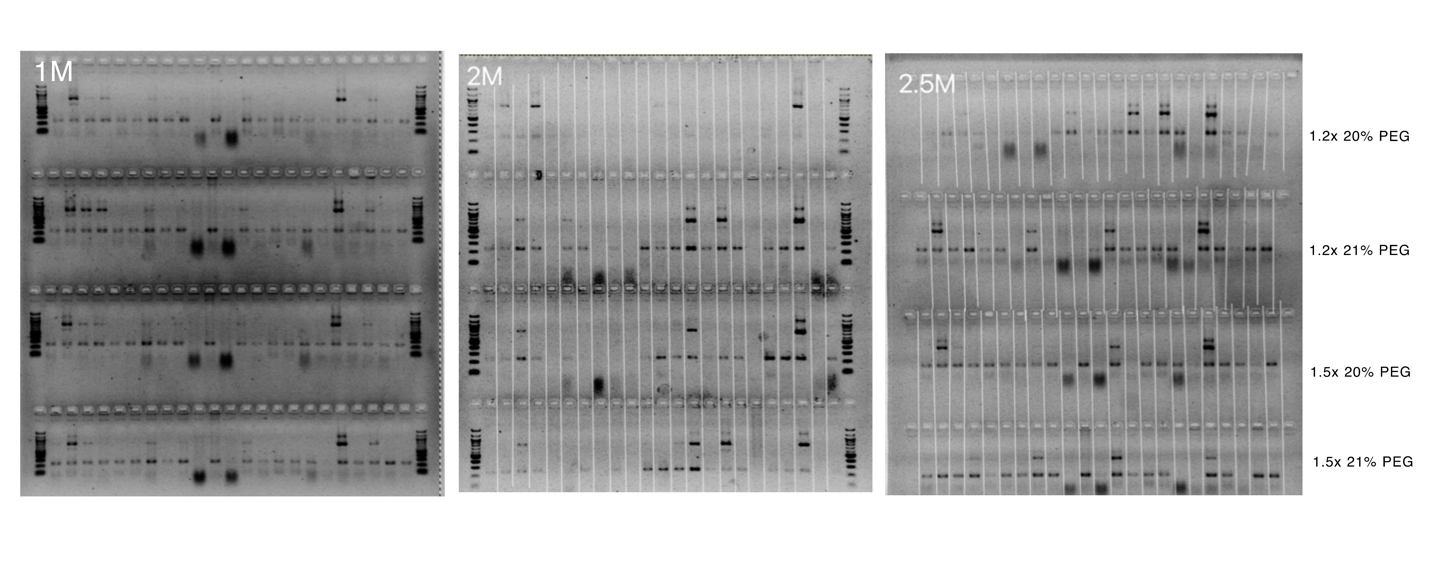

**Figure 8. Gel images from experiments #2, assessing amplification success given different concentrations of NaCl and PEG as well as different ratios of beads.** The number of bands observed as used as the success metric.

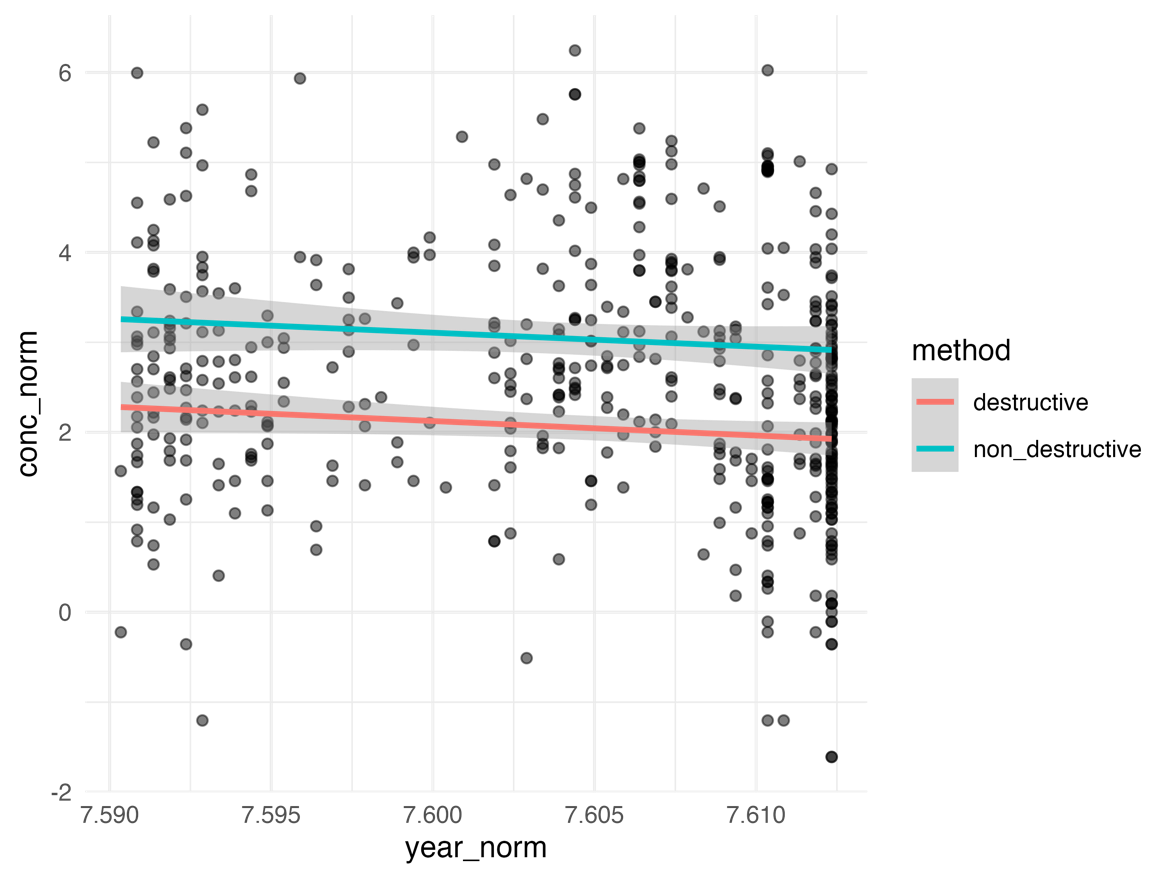

**Figure 9. Linear relationship between the DNA yields from non-destructive whole-body lysis and homogenized tissue.** There was no strong correlation between yield and age.
