## Supplemental Materials 3 for "Towards large-scale museomics projects: a cost-effective and high-throughput extraction method for obtaining historical DNA from museum insect specimens"

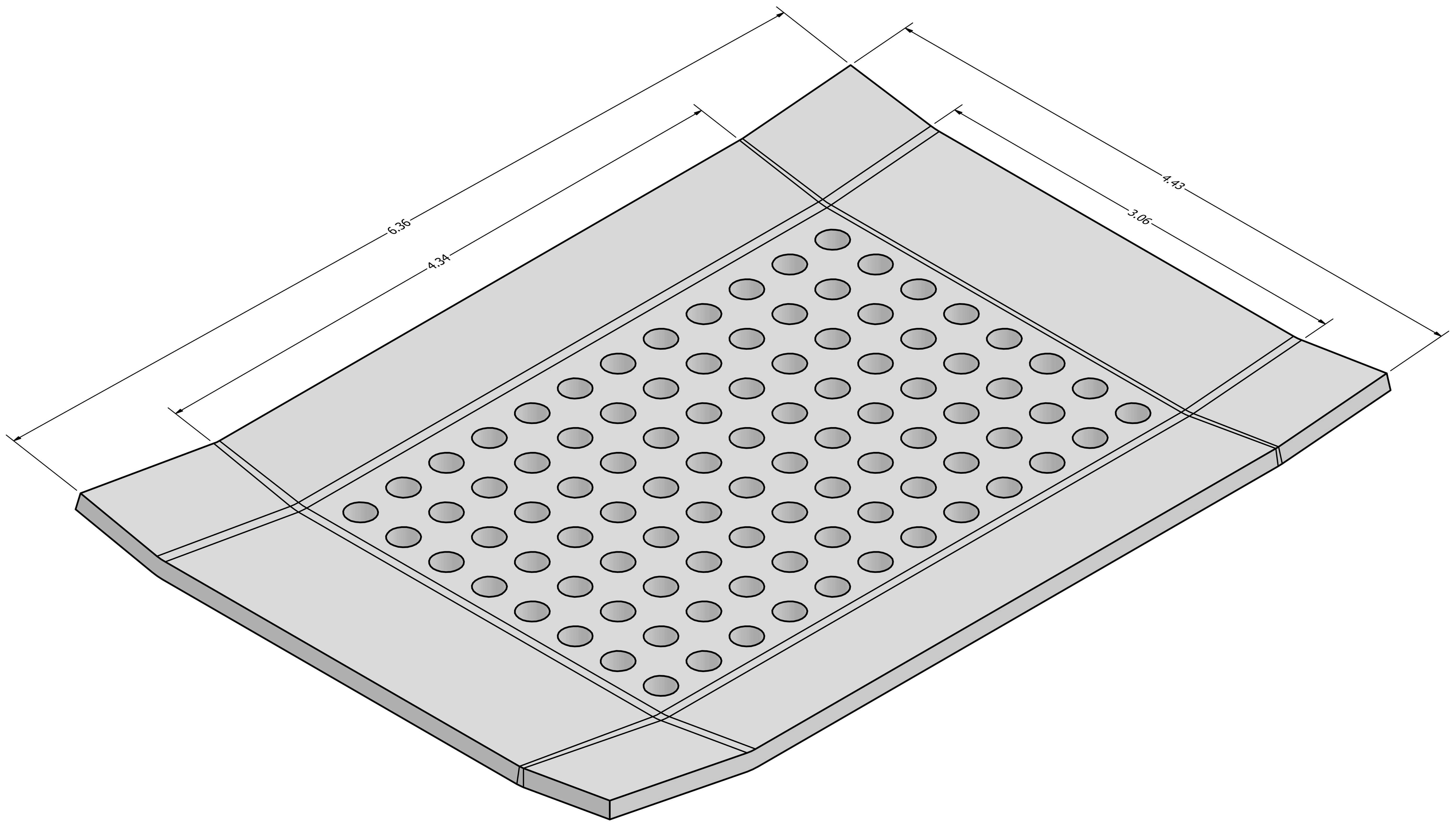


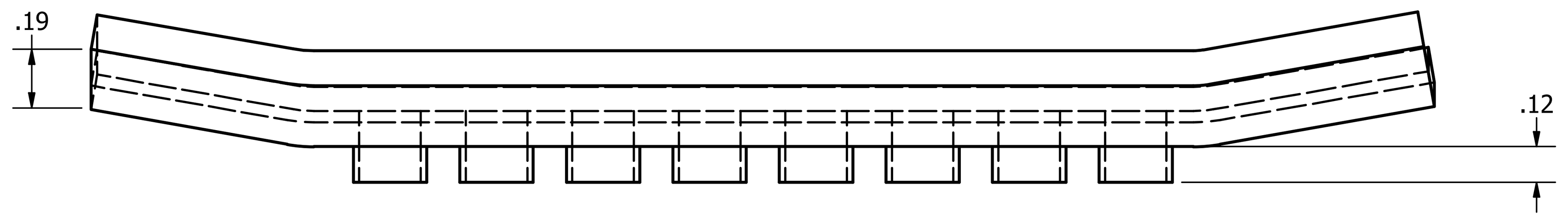


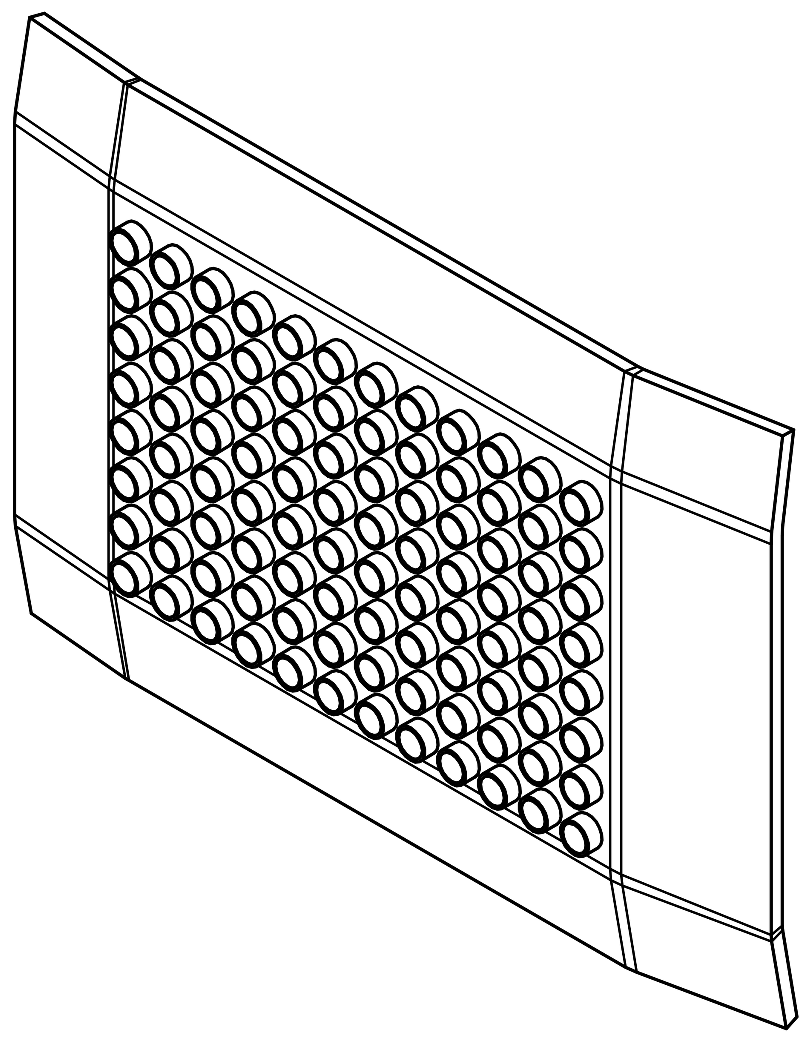

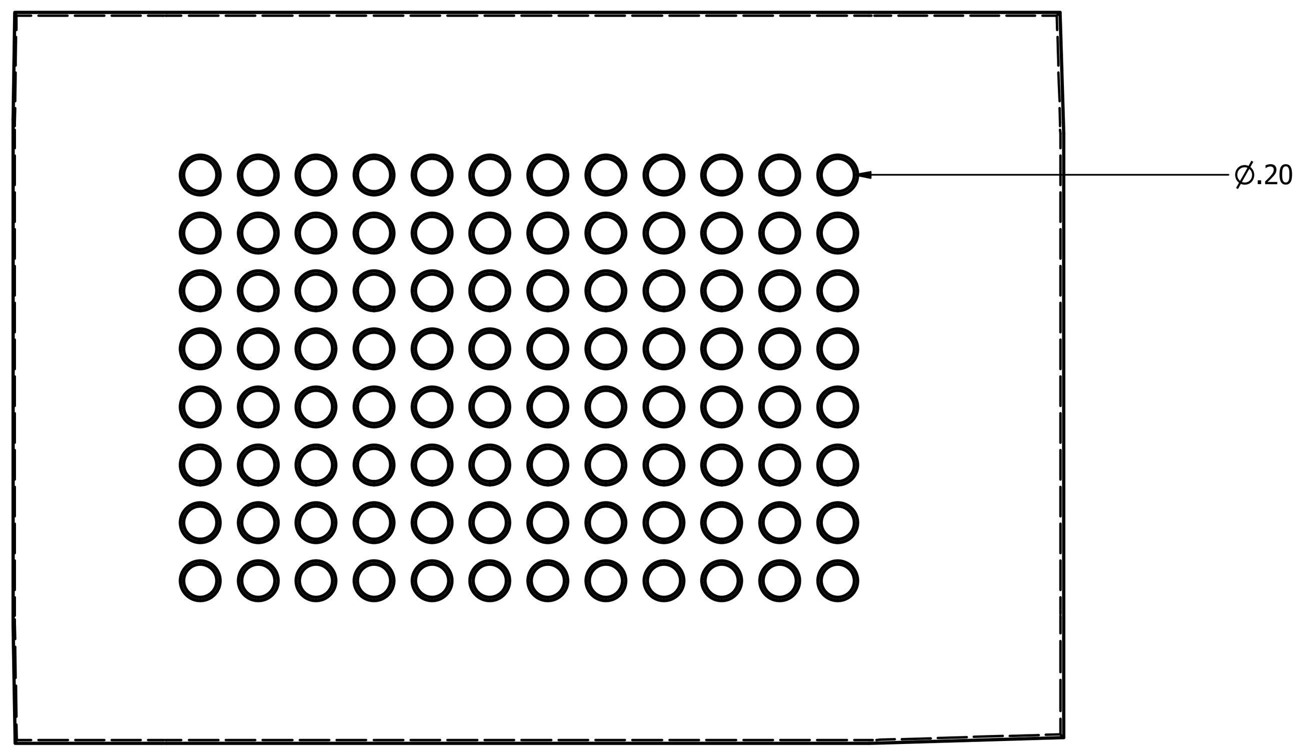


| | | |
|---|---|---|
| DRAWN tc | 6/18/2024 | CAS Experience Engineering for CAS Center for Comparative Genomics |
| CHECKED | | TITLE |
| QA | | ccgVial8x12 plate |
| MFG | | |
| APPROVED | | SIZE D / DWG NO / REV |
| | | SCALE 2 : 1 / SHEET 1 OF 1 |
