## Supplemental Materials 2 for "Towards large-scale museomics projects: a cost-effective and high-throughput extraction method for obtaining historical DNA from museum insect specimens"

**S2: SPRI Extraction Protocol**

1.0 **MATERIALS & EQUIPMENT**

**Reagents**

- Cell lysis buffer

*50mL 500mM EDTA*

*100mL 1M Tris-HCl*

*20mL 5M NaCl*

*50mL SDS (10%)*

*ddH2O to 1L*

- SPRI bead solution

*1mL Cytiva Sera-Mag SpeedBeads (Catalog: 09-981-123)*

*10.5g PEG-8000*

*10mL 5M NaCl*

*500uL 1M Tris-HCl*

*100uL 0.5M EDTA*

*27.5uL Tween*

*GeneRuler 1kb DNA Ladder (Catalog: SM0311)*

- GoldBio Proteinase K (Catalog: P-480-100) in 20mg/ml concentration
- Lab-grade 100% ethanol
- ddH_2_O

**Equipment**

- Silicon-Carbide sharp particles (Catalog: 11079101sc) for tissue
- 3.2mm stainless steel beads (Catalog: 11079132ss) for tissue
- 1/32^nd^ stainless steel teaspoon for tissue
- 96S super magnet plate (Catalog: 23A00H884)
- 2 PCR plates
- High-throughput tools if available (Tissuelyzer, BenchSmart, multichannel pipettes, repeater pipettes)

2.0 **PROCEDURES**

| **Specimen** | **Total volume** | **Proteinase K* (20mg/ml)** | **Cell lysis buffer** | **RPM** | **Time** | **Lysate volume** |
| --- | --- | --- | --- | --- | --- | --- |
| Tissue | 150uL | 2.25uL | 147.75uL | 180 | 24 hr | 50uL |
| Whole body  *small (soft)* | 150uL | 0.75uL | 149.25uL | 80 | 4 hr | 50uL |
| Whole body  *small (hard)* | 150uL | 2.25uL | 147.75uL | 180 | 16 hr | 50uL |
| Whole body  *medium (soft)* | 300uL | 1.5uL | 298.5uL | 80 | 16 hr | 50uL |
| Whole body  *medium (hard)* | 300uL | 4.5uL | 295.5uL | 180 | 24 hr | 25uL |
| Whole body  *large (soft)* | 600uL | 3uL | 597uL | 80 | 16 hr | 25uL |
| Whole body  *large (hard)* | 600uL | 9uL | 591uL | 180 | 24 hr | 25uL |

** Option to increase proteinase K concentration to 1mg/ml for small amounts of tissue or difficult tissue types; adjust volumes as necessary. We do not recommend higher concentrations as this can interfere with nucleic acid binding and separation of beads.*

**A. Lysis**

Tissue (destructive)

1. UV specimen plate for 5 mins with lids removed
2. Add sharp particles to each well using 1/32^nd^ teaspoon
   a. Optional: use plate attachment (S3, S4)
3. Add two steel beads to each well
   a. Optional: use plate attachment (S3, S4)
4. Make master mix (MM) of cell lysis buffer and proteinase K
   a. Refer to above table for volumes
5. Dispense 1**50uL** of MM to each well using repeater
6. Spin down plate
   a. Ensure tissue is submerged in buffer; push down with pipette tip if necessary
7. Tape kimwipe over capped columns, ensuring caps are completely secure
8. Place clear lid on plate
9. Place in tissuelyzer for 40 seconds at 20 Hz.
   a. Rotate plate and repeat
   b. Can use pestles for tissue homogenization instead
10. Spin down plate, ensuring no tissue is on lids
11. Place into the incubator at 55° for 24 hours (overnight) with shaker set to 180 RPM

Whole-body (non-destructive)

1. If samples were stored in ethanol, ensure no ethanol remains
2. UV specimen plate for 5 mins without lids
3. Make master mix (MM) of cell lysis buffer and proteinase K
   a. Refer to above table for volumes
4. Dispense MM in volume listed in the table for specimen type
5. Spin down plate
   a. Ensure specimen are completely submerged into buffer
6. Place into incubator at 55°
   a. Refer to above table for lysing time and RPM for shaking

**B. Bead Cleaning**

1. Add beads across a PCR plate with repeater pipette if available
   a. Add 37.5uL for 25uL of lysate or 75uL of beads for 50uL of lysate
2. Add lysate in the correct volume and mix by pipetting 10x
   a. Refer to table above for volumes
3. Transfer any remaining lysate to separate plates and freeze if desire for future
4. Centrifuge very briefly
   a. Beads should be fully homogenized and not settled
5. Incubate plate at room temperature for 10 minutes
6. While incubating, make 50mL of 80% ethanol with lab grade ethanol (40mL 100% and 10mL ddH2O) and put in freezer
   a. Make ethanol fresh every day as ethanol will absorb moisture from the air
7. Turn on incubator to 37 degrees
8. Move plate to magnet after incubation
   a. Make sure magnet and plate are sitting flat
9. Leave on magnet for 5 minutes or until liquid is completely clear
   a. Get EtOH from freezer at this stage and place in ice
10. Remove supernatant and discard into a waste container
    a. Perform using BenchSmart or multichannel pipette
11. Expel lysate tips into a tip box for later use

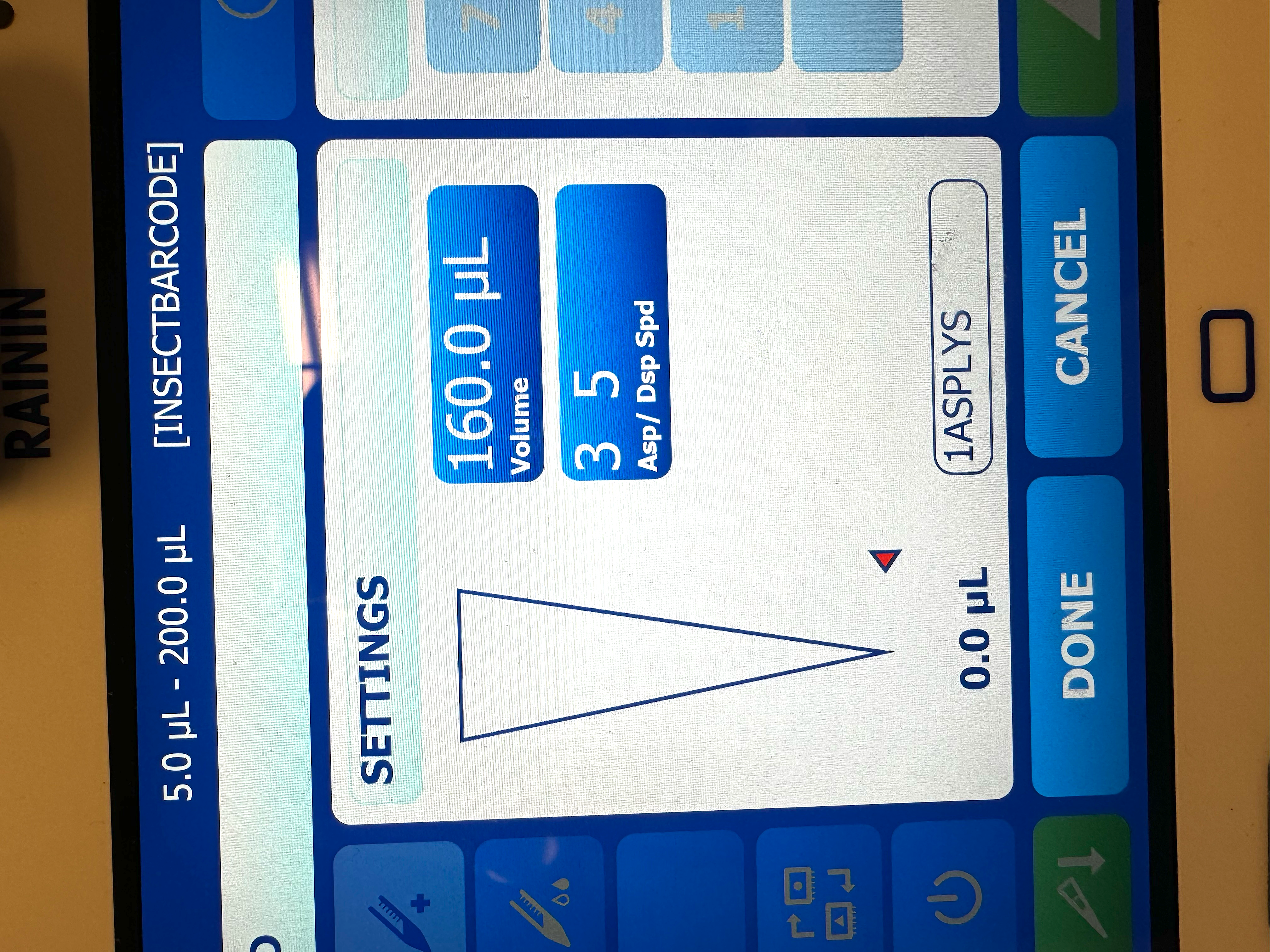

12. Add 150uL of cold ethanol to plate using either BenchSmart or multichannel pipette

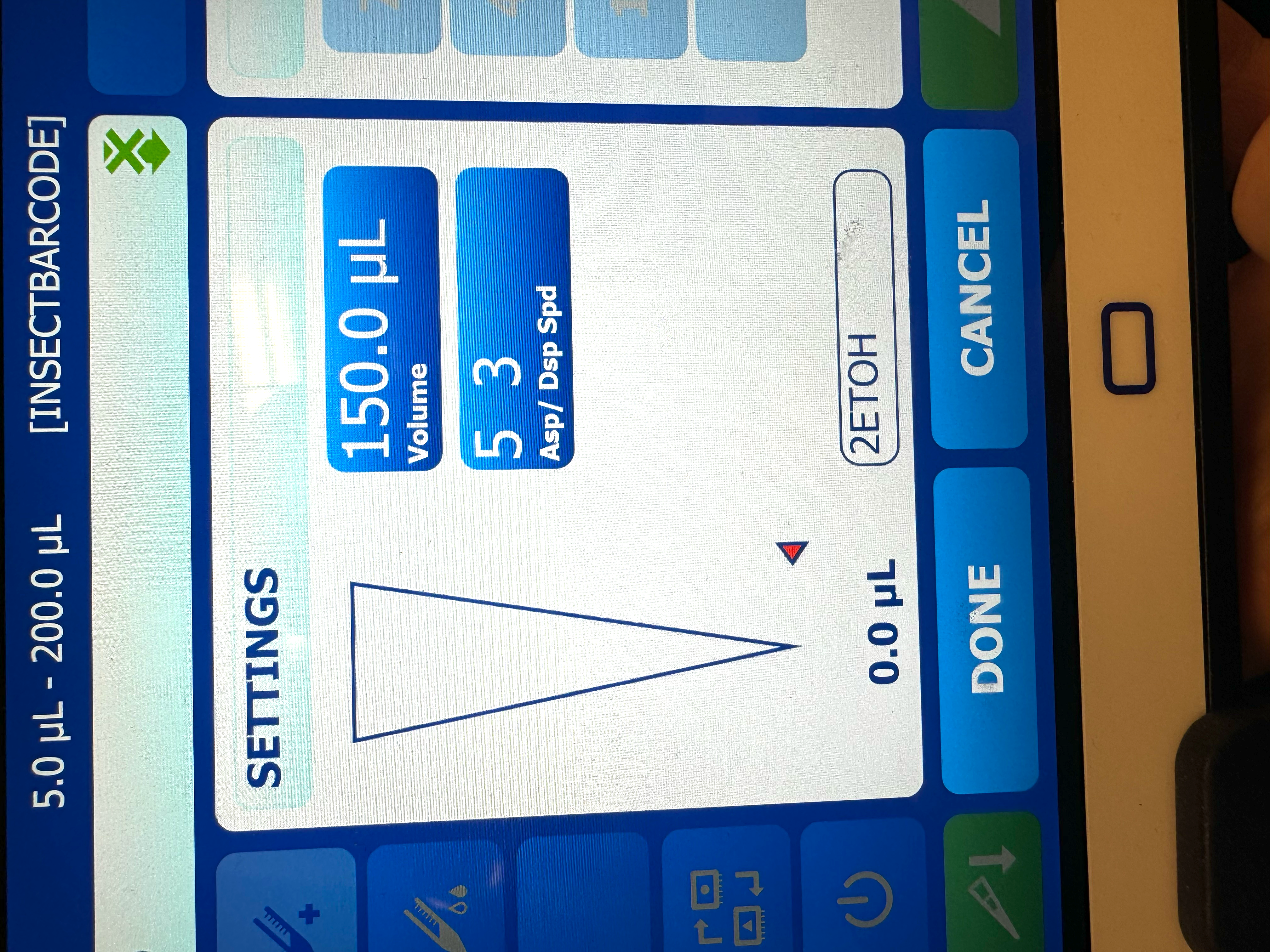

13. Mix EtOH up and down to wash beads x3 at a low speed
    a. Beads should not be disturbed

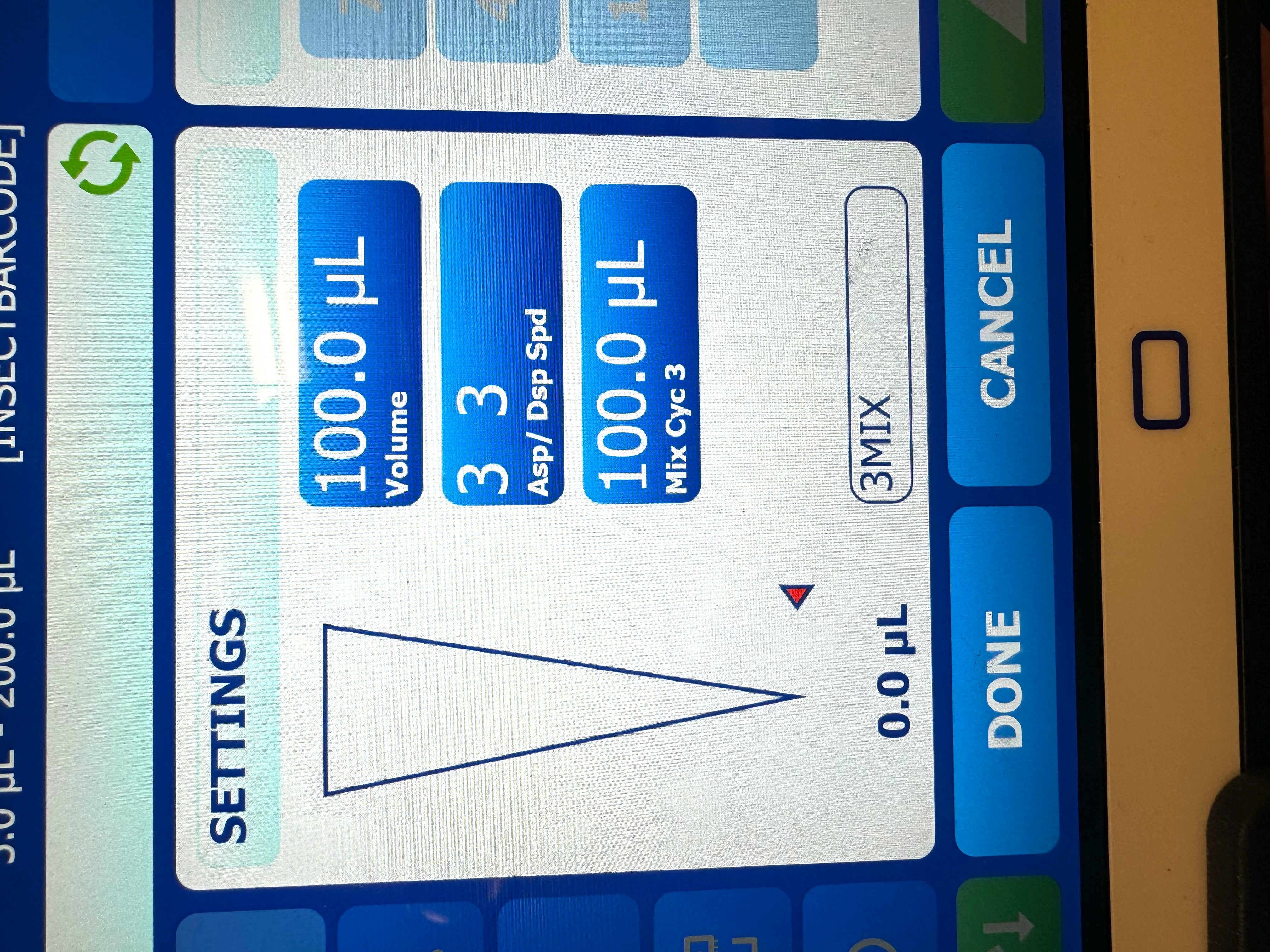

14. Incubate in ethanol for 1 minute
15. Add the lysate tips and remove ethanol

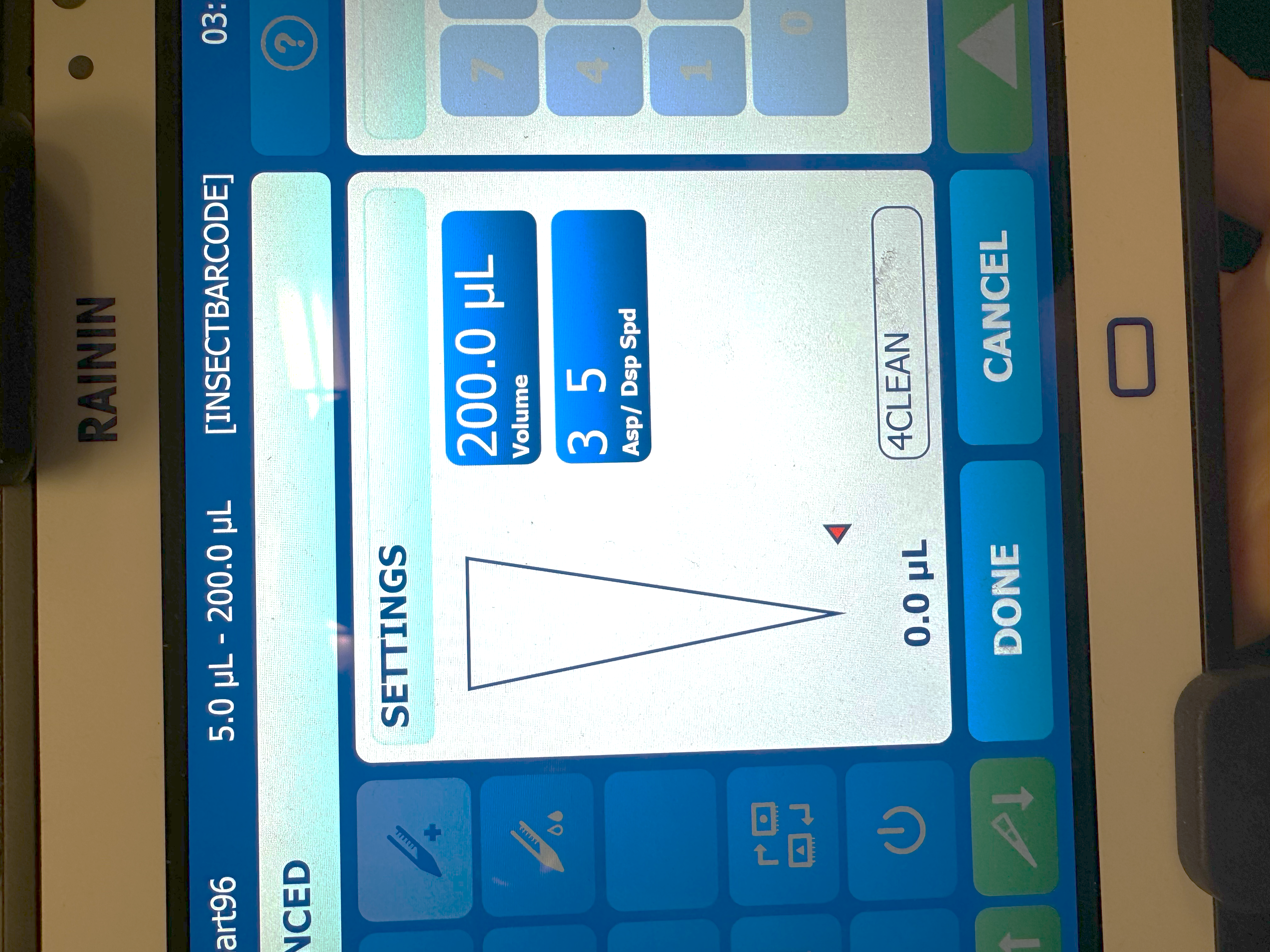

16. Repeat 10-15
    a. Remove ethanol entirely; use single channel to remove any remaining ethanol if necessary
17. Let any small amounts of ethanol evaporate for 2 minutes
    a. Do not overdry beads
18. Remove plate from magnet
19. Add 27uL of ddH_2_O to each well
    a. Use either benchsmart or repeater pipette

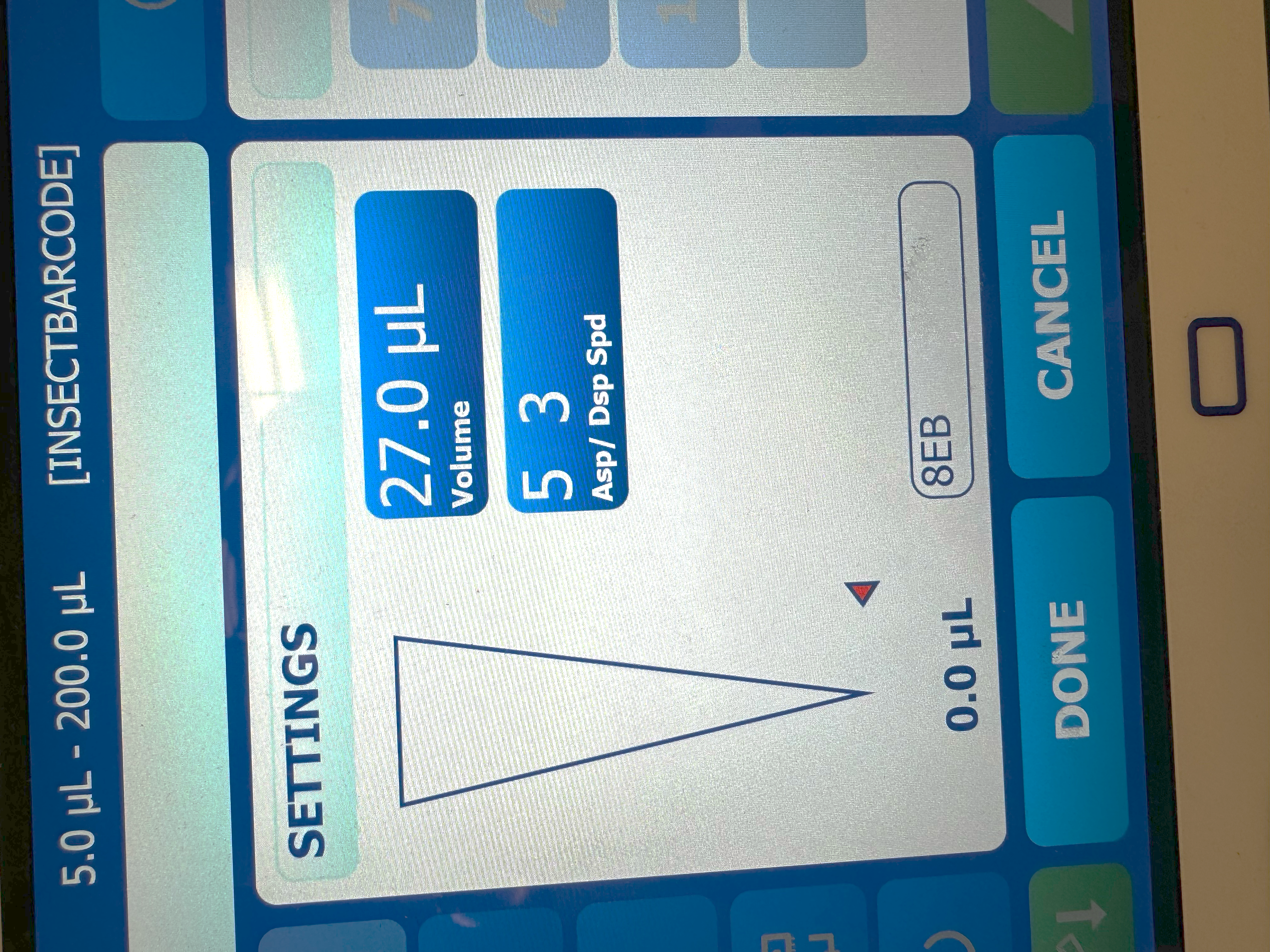

20. Add strip lids to plate and invert 10x
    a. Beads should be fully homogenized; flick plate if necessary
21. Briefly spin down
    a. Beads should remain homogenized
22. Incubate at 37 degrees for 5 minutes and another 5 minutes at room temperature
    a. Beads should be completely homogenized; flick or invert beads throughout process to ensure
23. Place plate back on magnet and leave for 2-5 minutes, until liquid is clear
24. Transfer 25uL of the eluted DNA to a new plate
25. Add strip lids
26. Discard lysate tips
3.0 **Recipe for solution: 1M 21% SPRI beads**
1. Make TE in 50 mL conical.
   a. TE = 500 uL 1 M Tris + 100 uL .5 M EDTA, fill to 50mL mark with ddH20
2. Vortex beads until fully homogenized, and transfer 1 mL to a 1.5 mL tube
3. Place tube on magnet stand.
4. When solution is clear, remove and discard supernatant with P200 pipette.
5. Add 1 mL TE to beads, dispensing directly against beads
6. Remove from magnet stand, vortex, and spin
7. Place on magnet stand
8. Repeat 4-7 two times
9. In a 50 mL conical, add 10.5g PEG-8000
10. Add 10mL 5 M NaCL
11. Add 500uL 1 M Tris-HCl
12. Add 100 uL .5M EDTA
13. Fill conical to ~49 mL mark with ddH20
14. Vortex and/or invert conical until PEG-8000 is fully dissolved and suspended into solution.
    a. Upon sitting, solution should be clear
15. Add 27.5 uL Tween 20, mix gently
16. Add 1 mL washed beads to solution
17. Fill to 50mL mark

*Ladder cleaning test*

18. Prepare fresh 80% ethanol
19. Make five tubes of 2uL of GeneRuler (must be this ladder) and 18uL ddH2O
20. To each tube, add a different ratio of beads:
    a. 0.8x = 16uL
    b. 1.2x = 24uL
    c. 1.5x = 30uL
    d. 1.8x = 36uL
    e. 2.1x = 42uL
21. Incubate for 5 minutes
22. Place on magnet stand and wait until solution is clear
23. Remove supernatent
24. Add 500uL of 80% ethanol
25. Incubate on magnet for another minute
26. Remove ethanol
27. Repeat steps 23-25 once
28. Let any remaining ethanol evaporate for 1-2 minutes
29. Rehydrate with 20uL of ddH2O, ensuring beads are fully homogenized
30. Incubate for 1 minute
31. Place on magnet
32. Transfer elute to individual tubes
33. Mix with loading dye provided with GeneRuler ladder
34. Run on gel
35. Store bead solution at 4 degrees wrapped in foil
